# Activation of two noncanonical R proteins by an insect effector confers plant immunity to aphid infestation

**DOI:** 10.1101/2024.07.06.601796

**Authors:** Kang Lei, Dong Tian, Yutao Shao, Faming Wang, Jinhua Chang, Si Nian Char, Guangwei Li, Zhenying Dong, Jianping Zhang, Jiang-Hui Cui, Songmin Zhao, Jingjing Li, Hua Liu, Guo-Qing Liu, Peng Lv, Mingshu Wei, Xiaohuan Jin, Qisheng Song, Bing Yang, Kunpu Zhang, Di Wu, Dao Wen Wang

## Abstract

Molecular characterization of resistance genes is crucial for efficiently understanding and fortifying plant immunity against insect herbivores. Here we report that RMES1A and RMES1B proteins confer resistance to the sorghum aphid *Melanaphis sorghi* when activated by an insect effector MsEF1. Map-based cloning plus genetic analysis of knockout mutants confirm that RMES1A and RMES1B are both required for aphid resistance. Upon aphid attack, RMES1A and RMES1B expression is elevated in the sclerenchyma cells and vascular bundles of leaves; the two proteins interact with MsEF1 in the exocysts, thus upregulating key defense processes such as reactive oxygen species burst. Structural modeling predicts that RMES1A and RMES1B each carry an ATP binding site and two leucine-rich-repeat domains but lack coiled-coil or Toll/Interleukin-1 receptor/resistance domain, thus likely representing a new type of resistance controlling proteins in plants. Our work reveals new genes and mechanisms for further deciphering and improving plant immunity to insect pests.

## INTRODUCTION

Aphids are among the most common insect pests that decrease worldwide crop productions.^1,2^ To efficiently breed aphid resistant crops, a solid understanding of the molecular basis of plant resistance to aphid herbivory is highly desirable.^1,3,4^ To date, two types of immunities, pattern-triggered immunity (PTI) and effector-triggered immunity (ETI), have been found in both plant-pathogen and plant-insect interactions.^5,6,7^ PTI offers a basal resistance and is activated after sensing pathogen or herbivore-associated molecular patterns by cell surface-located pattern recognition receptors (PRRs), whereas ETI confers a more robust race-specific resistance orchestrated after sensing pathogen or herbivore effectors by intracellularly located nucleotide-binding leucine-rich repeat receptors (NLRs). PRRs are usually receptor-like kinases (RLKs) or receptor-like proteins (RLPs), while NLRs have been divided into three types, coiled-coil (CC) NLRs (CNLs), Toll/Interleukin-1 receptor/Resistance (TIR) protein NLRs (TNLs), and RPW8-like CC domain (RPW8) NLRs (RNLs).^6^ Many PRRs require co-receptors during their regulation of PTI, and in many cases, both sensor and helper NLRs are needed for executing ETI.^6,7^

Among the 20 major-effect insect resistance proteins characterized by now,^3,7–15^ only three have been shown to promote host plant resistance after recognizing their cognate effectors from the feeding insects. A LRR-RLP conserved in Phaseoloid, plants has been demonstrated to initiate immune signaling and defense responses to herbivory caterpillars after recognizing the effector protein inceptin.^13,14^ An EDR1-like protein kinase has been shown to promote tobacco defense against the aphid *Myzus persicae* via interacting with the effector CathB3.^9^ Recently, the rice brown planthopper (BPH) resistance protein BPH14, a canonical CNL, is reported to confer ETI to BPH after binding the effector protein BISP.^16^ Nevertheless, the avirulence effectors and activation mechanisms remain poorly understood for many of the reported insect resistance proteins, including 10 CNLs from rice (BPH1, 2, 7, 9, 10, 18, 21, and 26), tomato (Mi-1), and melon (Vat) and two rice LRR proteins (BPH6 and BPH30).^8,12,17,18^

Sorghum is a highly valuable food, feed, and energy crop, but its stable production is seriously threatened by the aphid *Melanaphis sorghi* (*MES* hereafter), especially by a highly aggressive *MES* clone that is spreading rapidly in many American countries and the world.^19–22^ Consequently, there is now a wide interest in identifying the genes functioning in sorghum resistance to *MES* for breeding resistant cultivars.^19,22–30^ However, no definite PRRs or NLRs that function in sorghum resistance to *MES* have yet been molecularly characterized. Previously, we mapped *RMES1* locus, conferring a strong resistance MES, to a 126 kb genomic region on the short arm of chromosome 6 using the genetic populations developed with HN16 (an elite *MES* resistant grain sorghum variety) and BTx623 (susceptible to *MES* and with available genome sequence) as parents.^31^ Here we report that two new genes (designated as *RMES1A* and *RMES1B*) within the *RMES1* locus are both required for sorghum resistance to *MES*. We further identify a *MES* effector protein (namely MsEF1) that binds to RMES1A and RMES1B and strengthens their interaction in the exocysts, which leads to immune signaling and upregulation of defense responses (e.g., ROS burst and callose deposition) to *MES* feeding. RMES1A and RMES1B are more closely related to the rice resistance proteins BPH6 and BPH30 than to typical CNLs and TNLs. They are predicted to carry an ATP binding site and two LRR domains but lack the coiled-coil or Toll/Interleukin-1 receptor/resistance domain. We also analyzed natural variations of *RMES1A* and *RMES1B* in 166 worldwide sorghum germplasm lines, which not only validates the function of *RMES1A* and *RMES1B* but also enriches the germplasm with functionally tested *RMES1* locus for efficient molecular breeding of *MES* resistant sorghum in the future.

## RESULTS

### Map-based cloning and functional validation of *RMES1* locus

Based on our previous work,^31^ we further mapped *RMES1* to a 25 kb region using newly developed DNA markers and 5,136 BC5F2 plants segregating for *RMES1* (Figure 1A). Although flanked by two genes (named as *LRRP2* and *LRRP3*, respectively) predicted to encode LRR containing proteins (Figure 1A), the 25 kb region did not carry any protein-coding genes in the genome assembly of BTx623, indicating probable chromosomal structural variation at this location among different sorghum genotypes. Hence, we constructed a bacterial artificial chromosome (BAC) library for the *MES* resistant line HN16 and obtained 194 kb genomic sequence through sequencing four overlapping BAC clones that were identified using the DNA markers flanking *RMES1*. Subsequent syntenic genomic analysis revealed that the marker-tagged 25 kb region in BTx623 was expanded to 169 kb in HN16, which harbored two additional *LRR* genes (*RMES1A* and *RMES1B*, Figure 1A) not present in BTx623. Thus, they were considered as candidate causal genes of *RMES1*. The open reading frames of *RMES1A* and *RMES1B* were devoid of introns, and their coding region was all 3,099 bp (including stop codon). Their deduced proteins, both having 1032 amino acids and a predicted molecular mass of approximately 118 kDa, were 96.2% identical. Notably, *RMES1A* and *RMES1B* were also highly similar to their flanking genes (*LRRP1*, *LRRP*2, and *LRRP3*), with their deduced proteins being 62.0%-82.9% identical.

**Figure 1.**
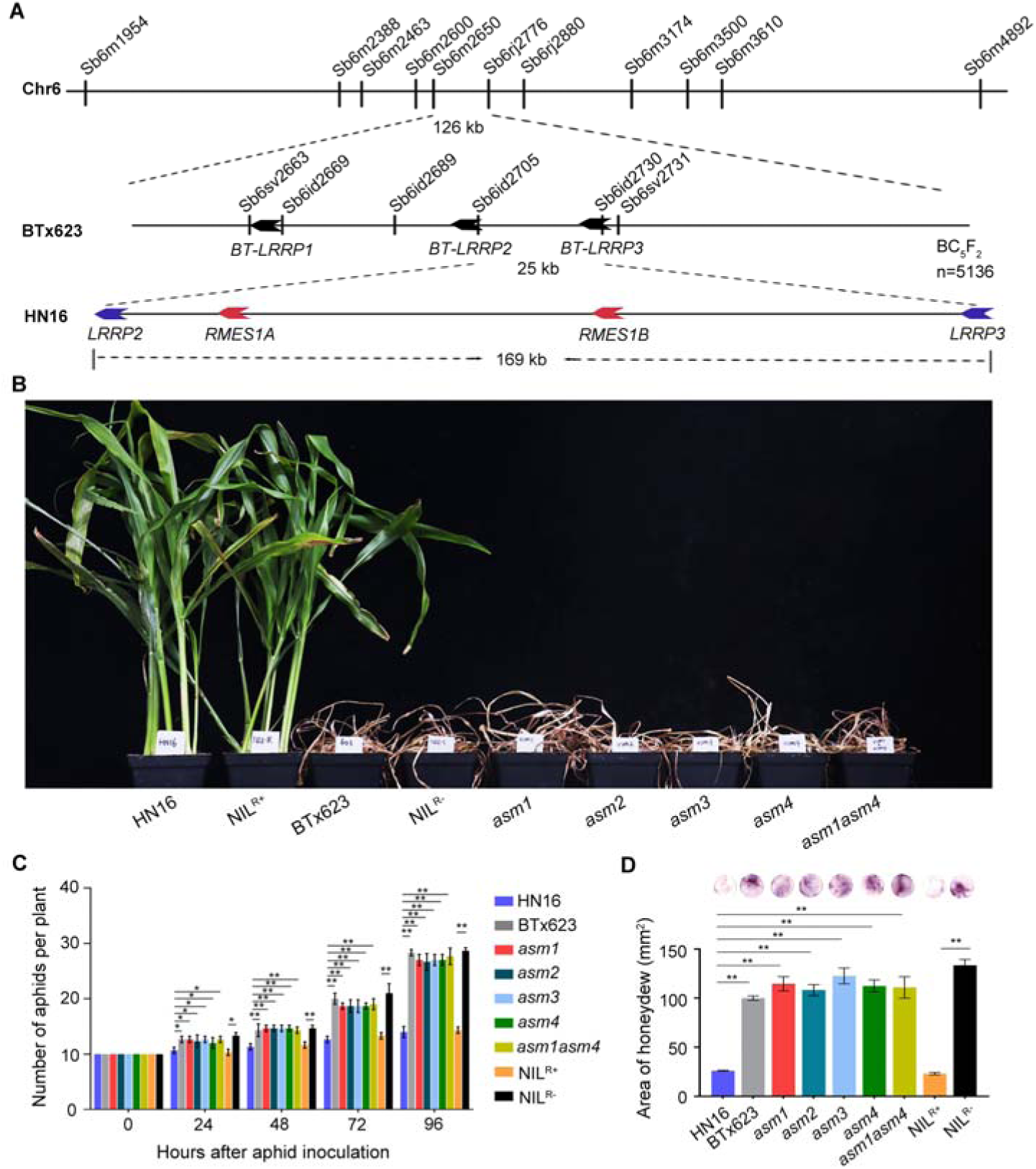
Map-based cloning and functional validation of *RMES1* genes. (A) *RMES1*, previously mapped on sorghum chromosome 6 between the markers Sb6m2650 and Sb6rj2776,^31^ was further localized to a 25 kb region using 5,136 BC_5_F_2_ plants according to the genome sequence of the *MES* susceptible line BTx623. This region, flanked by *LRRP1*, *LRRP2*, and *LRRP3*, contained no genes in BTx623. However, it was expanded to 169 kb in the *MES* resistant cultivar Henong 16 (HN16) and contained two additional genes (*RMES1A* and *RMES1B*). (B) Mutation of *RMES1A* and *RMES1B* leads to *MES* susceptibility. Five mutants with mutations in *RMES1A* (*asm1*, *asm2*), *RMES1B* (*asm3, asm4*) or both (*asm1asm4*) resembled BTx623 and NIL^R-^ in being susceptible to *MES*, whereas HN16 and NIL^R+^ exhibited resistance to *MES*. NIL^R+^ and NIL^R-^ were isogenic lines with or without *RMES1* locus. (C) The number of *MES* aphids in nine resistant or susceptible sorghum genotypes at different hours post inoculation. Error bars, means ± SE (n = 3). *p < 0.05; **p < 0.01 (Student’s *t* test, compared to HN16). (D) The amount of honeydew excreted by *MES* aphids on resistant or susceptible sorghum plants. The honeydew excreted on filter paper was stained using ninhydrin. Both the area and intensity of honeydew staining were recorded. Error bars, means ± SE (n = 3). **p < 0.01 (Student’s *t* test, compared to HN16).

Next, we developed four *MES* susceptible mutants (*asm1*, *asm2*, *asm3*, and *asm4*) using HN16. While *asm1* and *asm2* were prepared by mutagenesis with diepoxybutane or ethyl methane sulfonate, *asm3* and *asm4* were identified from two independent fast neutron mutant families, respectively. Both *asm1* and *asm2* carried nucleotide deletions in *RMES1A* coding sequence, with the deduced asm1 and asm2 proteins lacking 59 residues or being prematurely truncated (Figure S1A). Genome PCR and genomic resequencing analyses showed that *asm3* and *asm4* lacked the entire *RMES1B* gene, with *RMES1A* remaining intact (Figures S1B and S1C). Besides, a double mutant (*asm1asm4*) lacking *RMES1A* and *RMES1B* was prepared by crossing *asm1* and *asm4*; two near isogenic lines with or without the *RMES1* locus, designated as NIL^R+^ and NIL^R-^, respectively, were selected from a BC7F2 family developed with BTx623 as recurrent parent (Figure S1D). In repeated *MES* feeding assays, BTx623, NIL^R-^, and the five mutants, but not HN16 and NIL^R+^, became wilted and dying after 10 days post inoculation (dpi) (Figure 1B). The number of inoculated aphids increased by more than 2.5 folds in BTx623, NIL^R-^, and the five mutants, whereas it was merely 1.1 folds in HN16 and NIL^R+^ by 96 hours post inoculation (hpi) (Figure 1C). Consistently, aphid honeydew was produced in significantly larger amounts in the seven susceptible genotypes than in HN16 and NIL^R+^ at 48 hpi (Figure 1D).

We further explored natural variations of *RMES1A* and *RMES1B* in 166 sorghum accessions collected from different countries (Table S1). We designated *RMES1A* and *RMES1B* sequences isolated from HN16 as wild type genes, the null alleles of *RMES1A* and *RMES1B* as *rmes1a-1* and *rmes1b-1*, respectively, and the variants showing amino acid substitutions in deduced RMES1A and RMES1B proteins as *rmes1a-2* and *rmes1b-2*, respectively. Based on the result obtained, the 166 lines were divided into five types (Table S1). Type I (n = 36) had both *RMES1A* and *RMES1B* genes, and all showed resistance to *MES*, whereas Type II (n = 115) carried *rmes1a-1* and *rmes1b-1* and exhibited aphid susceptibility. Type III (n = 9) carried *RMES1B* but the mutant *rmes1a-2* allele whose deduced protein showed amino acid substitutions mainly in two different segments of RMES1A, i.e., S1 (169-176) and S2 (194-197) (Table S1; Figure S2). Type IV (n = 5) harbored *rmes1a-2* and *rmes1b-1*, while Type V (n = 1) possessed *rmes1a-1* and *RMES1B* (Table S1). The latter three types of lines were all susceptible to *MES*, despite the presence of *RMES1B* in Types III and V materials (Table S1).

Lastly, CRISPR-induced knockout mutant was generated using the sorghum accession P89002 (PI 561847), which carried both *RMES1A* and *RMES1B* and exhibited resistance to *MES* infestation. A CRISPR mutant (*rmes1a1b-#25*), harboring knockout mutations in both *RMES1A* and *RMES1B*, was highly susceptible to *MES* feeding (Figure S3A-D), further corroborating the findings made by using the single and double *asm* mutants (Figures 1B and 1C). Taken together, the above genetic data strongly suggest that *RMES1A* and *RMES1B* are both required for sorghum resistance to *MES*.

### Impairment of key defense responses in *asm* mutants

We examined whether the *asm* mutants may be impaired in antixenosis and antibiosis, which deters aphid feeding and inhibits aphid reproduction, respectively.^32^ In free choice tests, the number of *MES* adults settled on the resistant genotypes (HN16 and NIL^R+^) generally decreased and remained low, whereas the converse was observed on the susceptible lines (*asm1*-*asm4*, *ams1asm4*, NIL^R-^, and BTx623) (Figure S4A). In no-choice tests, the nymphs newly reproduced by *MES* increased dramatically on all seven susceptible lines, with no or very few nymphs found on the two resistant genotypes (Figure S4B). Thus, both antixenosis and antibiosis function in *RMES1A* and *RMES1B* mediated resistance to *MES*.

Histochemical staining of *MES* inoculated sorghum leaves with diaminobenzidine (DAB), which efficiently detects localized hydrogen peroxide (H2O2) production in plant cells,^33^ showed increasingly intense H2O2 accumulation in more *MES* feeding sites in HN16 and NIL^R+^ (Figure 2A). In contrast, H2O2 accumulation occurred in fewer *MES* feeding sites with relatively low intensity in the seven susceptible lines (Figure 2A). H2O2 content was significantly elevated upon *MES* inoculation in HN16 and NIL^R+^, especially at 48 hpi, but this increase was not obvious and exhibited inconsistent fluctuations in the seven susceptible genotypes (Figure 2B). This result agrees well with the suggestion that elevated H2O2 accumulation is closely associated with the *MES* resistance exhibited by RTx2783 that carries *RMES1* locus.^29,34^ Callose deposition was also significantly upregulated in HN16 and NIL^R+^ but not the seven susceptible lines after *MES* inoculation, as uncovered by histochemical staining with aniline blue and quantitative measurement (Figure S5).

**Figure 2.**
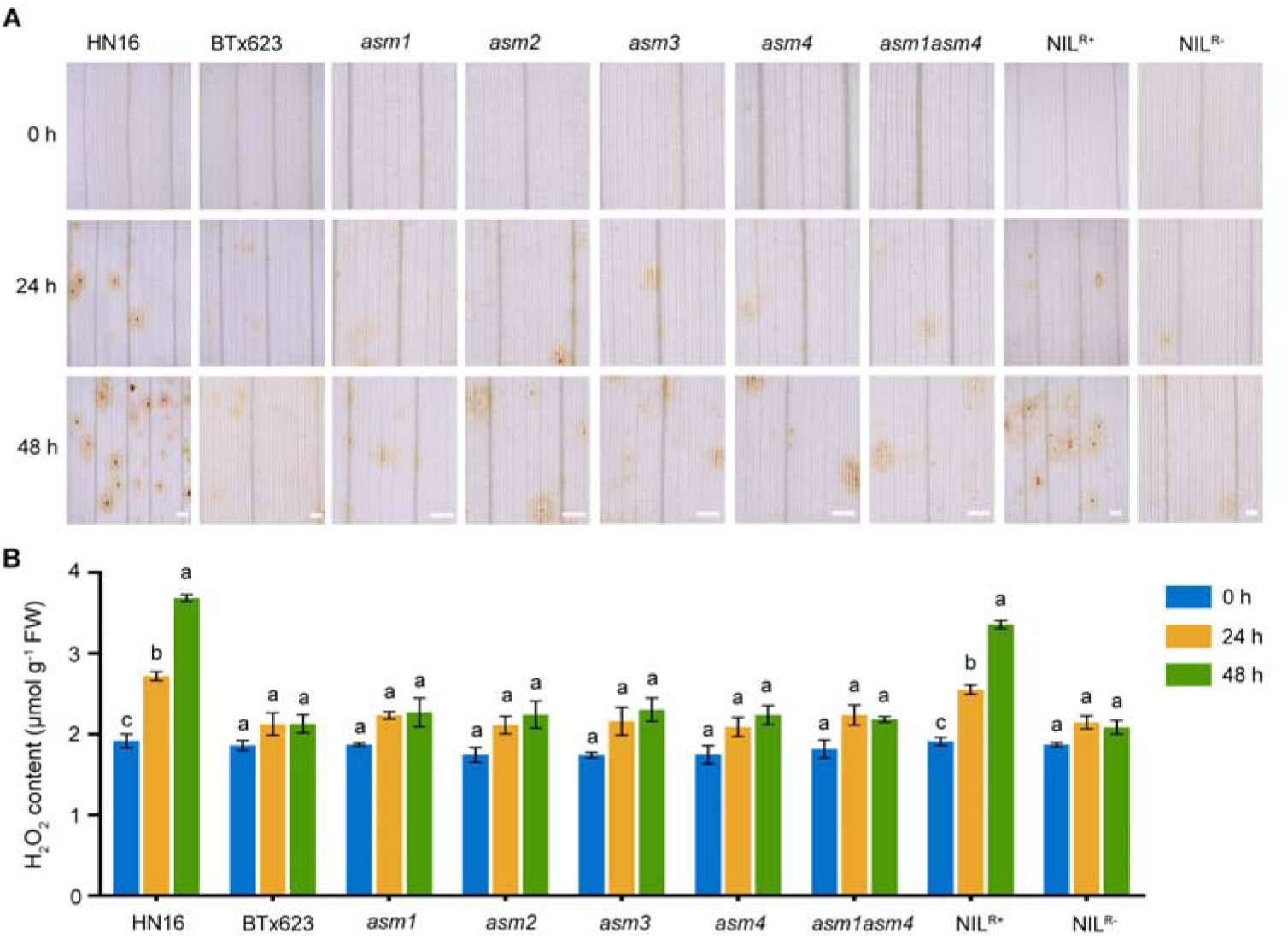
*REMS1A* and *RMES1B* are required for efficient induction of H_2_O_2_ burst upon *MES* feeding. (A) Detection of H_2_O_2_ accumulation in *MES* feeding sites by DAB staining in nine resistant or susceptible sorghum genotypes at 0, 24, and 48 h post inoculation (from top to bottom). Scale bars, 20 µm. (B) Quantitative comparison of H_2_O_2_ contents of sorghum leaves infested by aphids for 0, 24 and 48 h. Error bars, means ± SE (n = 3). Different letters above the histograms indicate significant differences by Tukey’s HSD test (p < 0.05).

### Expression and subcellular targeting of RMES1A and RMES1B

During the development of HN16 plants, *RMES1A* and *RMES1B* expression was detected in both vegetative (root, stem, and leaf) and reproductive (spike and seed) organs by RT-qPCR assays, with the expression level of *RMES1A* being significantly higher than that of *RMES1B*, especially in the stem and leaf tissues (Figure 3A). In the 14-day old HN16 seedlings inoculated with *MES*, *RMES1A* expression was highly upregulated with the peak detected at 24 hpi; *RMES1B* was similarly induced by *MES* although its expression levels were generally and significantly lower than those of *RMES1A* at the time points examined in this work (Figure 3B).

**Figure 3.**
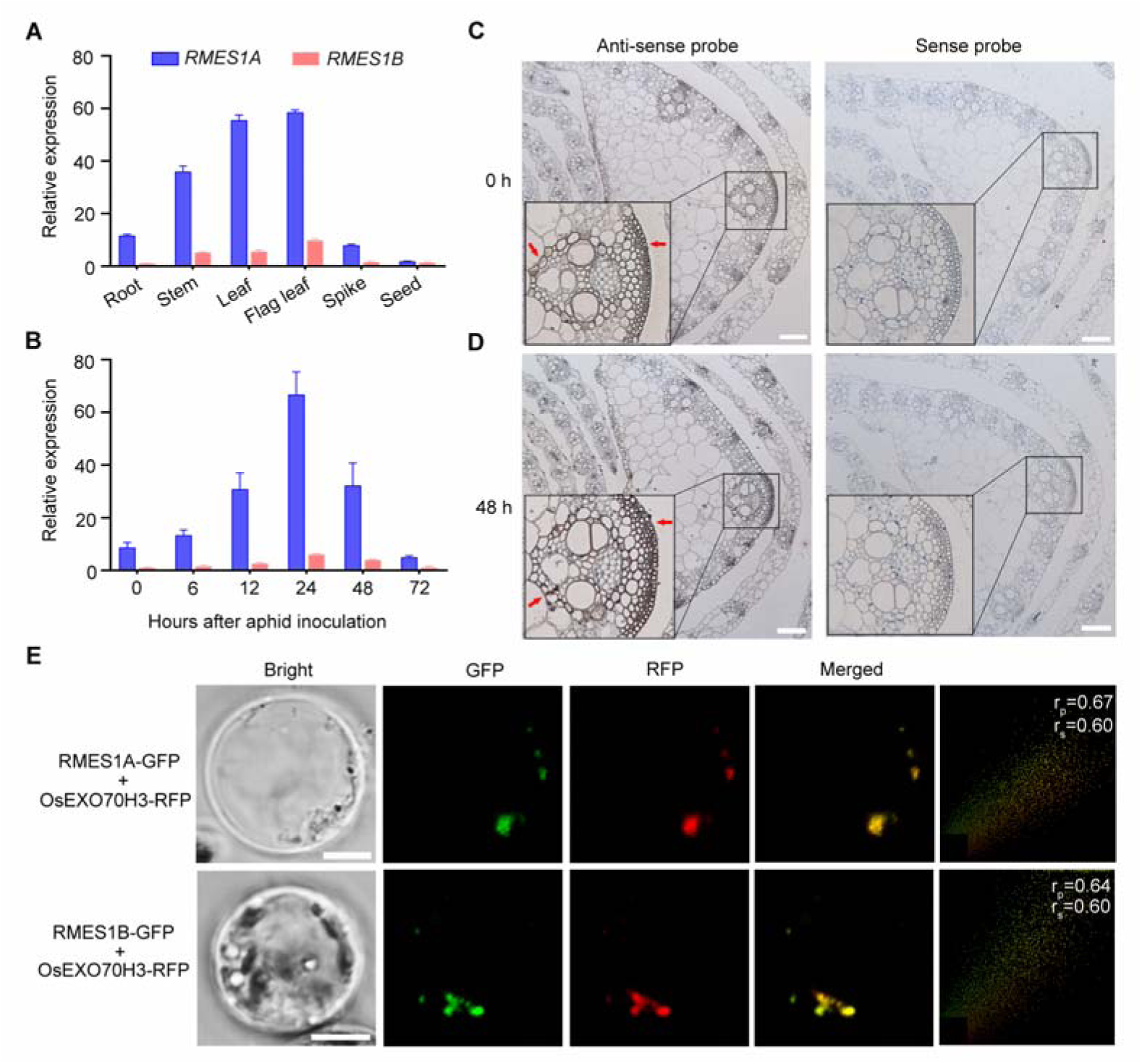
Expression and subcellular localization of RMES1A and RMES1B. (A) Quantification of *RMES1A* and *RMES1B* expression in various organs of HN16 plants without *MES* infestation. Error bars, means ± SE (n = 3). (B) Expression profiles of *RMES1A* and *RMES1B* at different hours post *MES* inoculation. Error bars, means ± SE (n = 3). (C, D) *In situ* hybridization of *RMES1A* and *RMES1B* mRNAs in HN16 leaf tissues after *MES* feeding for 0 (C) and 48 h (D). Hybridization with an antisense probe revealed strong signals in the sclerenchyma cells and vascular bundles (red arrows). Such signals were not detected in the negative controls performed with a sense probe. Scale bars,100 μm. (E) RMES1A and RMES1B are localized in sorghum exocysts. GFP-tagged RMES1A or RMES1B was co-expressed with an RFP-tagged rice exocyst subunit OsEXO70H3 in sorghum protoplasts. Overlapping of GFP and RFP fluorescence signals suggests co-localization of RMES1A and RMES1B with OsEXO70H3 in the exocysts. Pearson correlation coefficient (r_P_) and Spearman correlation coefficient (r_S_) are both in the range of +1 to-1, which indicate strong positive and negative correlations, respectively. Scale bars, 5 μm.

To investigate the spatial expression pattern of *RMES1A* and *RMES1B* before and after *MES* inoculation, we conducted RNA *in situ* hybridization assays with an antisense probe corresponding to the transcripts of both genes due to high nucleotide similarity (97.8%) in their coding sequence. *RMES1A* and *RMES1B* transcripts were detected in the sclerenchyma cells and vascular bundles of leaf sheaths, which were substantially elevated at 48 hpi of *MES* (Figures 3C and 3D). The spatial expression feature of *RMES1A* and *RMES1B* resembled that of *BPH6* and *BPH30*, both of which confer resistance to the suckling insect BPH in rice.^12,35^

The subcellular targeting of RMES1A and RMES1B was analyzed by expressing their C-terminal GFP fusion proteins in sorghum protoplasts. Confocal microscopy revealed the appearance of discrete green fluorescence structures in the cytoplasm after expressing RMES1A-GFP or RMES1B-GFP, which overlapped with the red fluorescence structures formed by the RFP fusion protein of OsEXO70H3 (Figure 3E), Because OsEXO70H3 is a subunit of the exocyst complex functioning in intracellular protein trafficking,^36^ these data suggest that RMES1A and RMES1B are located in the exocysts in sorghum cells.

### Interaction of RMES1A and RMES1B in the exocysts

The findings described above prompted us to test whether the two proteins may interact with each other or not. Indeed, RMES1A was found to interact with RMES1B in not only yeast two hybrid (Y2H) assays but also the split luciferase complementation (SLC) experiments conducted in tobacco leaves (Figures 4A and 4B). This interaction was validated by co-immunoprecipitation (Co-IP) and bimolecular fluorescence complementation (BiFC) assays in sorghum protoplasts (Figures 4C and 4D). Note that the BiFC signals resulted from RMES1A and RMES1B interaction overlapped with the red fluorescence of OsEXO70H3-RFP (Figure 4D), indicating that RMES1A and RMES1B interact in the exocysts. Furthermore, we found that the asm1 mutant protein, which had an internal deletion of 59 residues in the C-terminal part of RMES1A (Figure S1A), failed to interact with RMES1B in either yeast cells or sorghum protoplasts (Figures 4C and 4D). These protein-protein interaction data, plus the findings that *MES* resistance was lost in the *asm1* mutant (Figure 1B) and the sorghum lines with normal RMES1B but mutant RMES1A (Table S1), support the crucial importance of RMES1A and RMES1B interaction in conferring *MES* resistance.

**Figure 4.**
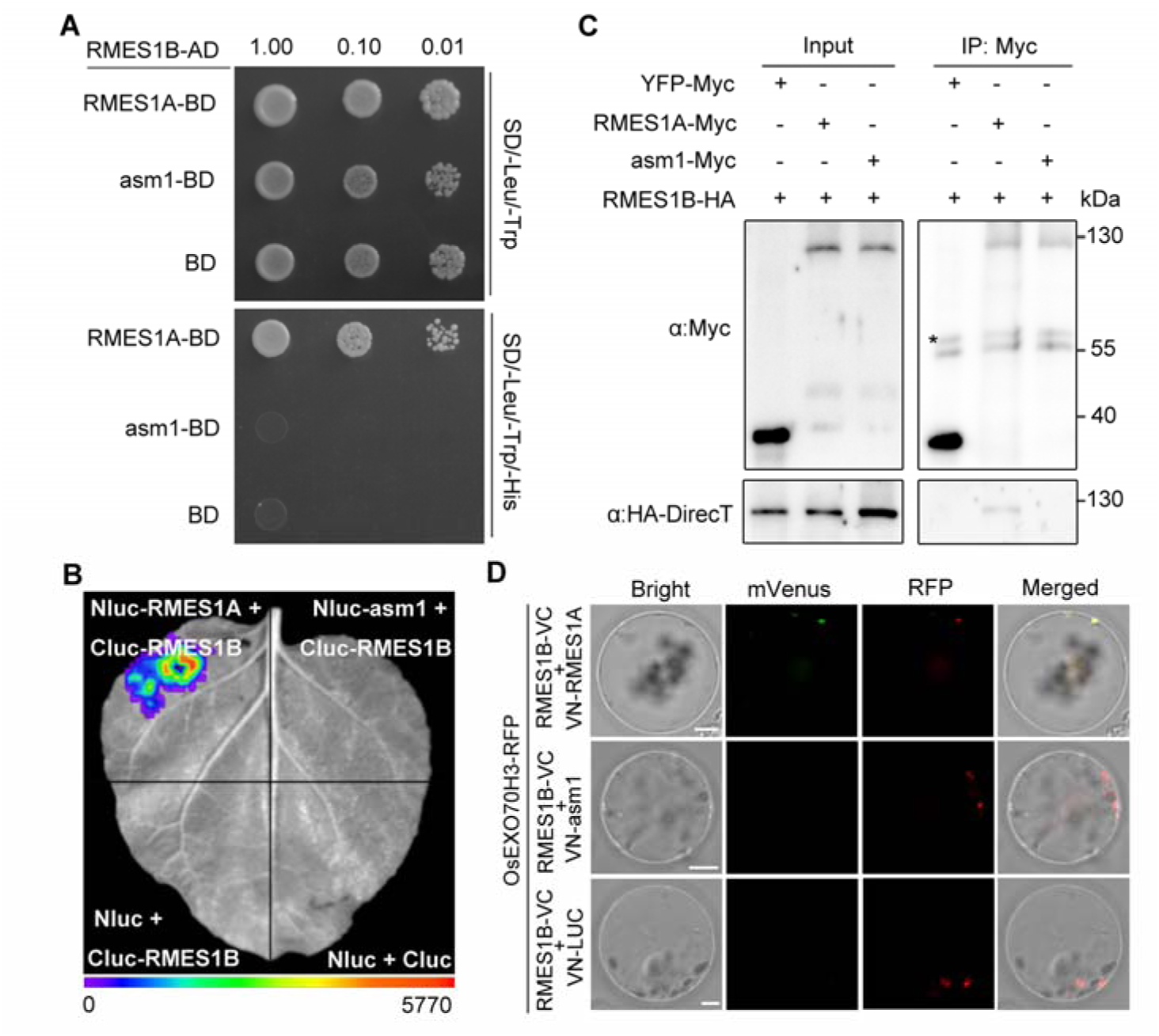
RMES1A and RMES1B interact in sorghum exocysts. (A-C) Positive interaction between RMES1A and RMES1B detected using yeast two hybrid (Y2H) (A), split luciferase complementation (SLC) (B), and co-immunoprecipitation (Co-IP) assays (C). The asm1 mutant failed to interact with RMES1B in these assays. In Y2H, RMES1B was fused with activation domain (AD), while RMES1A or asm1 was fused with DNA-binding domain (BD). The recombinant yeast cultures were serially diluted, with yeast colony growth on the SD/-Leu/-Trp/-His medium indicating occurrence of protein-protein interaction. In SLC, RMES1A or asm1 was fused to N-terminus of luciferase (Nluc), and RMES1B was fused to C-terminus of luciferase (Cluc). Tobacco leaf was co-infiltrated with the GV3101 strains carrying different pairs of constructs. Only the upper left sector showed positive interaction signals. In Co-IP, RMES1B-HA was co-expressed with RMES1A-Myc, asm1-Myc or YFP-Myc in sorghum protoplasts. Total proteins were immunoprecipitated using anti-Myc antibody, followed by immunoblotting with anti-HA-DirecT and anti-Myc antibodies, respectively. YFP-Myc was used as a negative control. Asterisk indicates the bands of IgG heavy chain. (D) RMES1A and RMES1B interact in sorghum exocysts revealed by bimolecular fluorescence complementation (BiFC) assays. RMES1B was fused with C-terminus of mVenus (VC), while RMES1A or asm1 was fused with N-terminus of mVenus (VN). VN fused luciferase (VN-LUC) served as a negative control. The different pairs of constructs, together with the one specifying OsEXO70H3-RFP, were transiently expressed in protoplasts. Yellow fluorescence signals indicate reconstituted mVenus through RMES1A-RMES1B interaction. Red fluorescence signals caused by OsEXO70H3-RFP mark the exocysts. Scale bars, 5 μm.

### Activation of *RMES1* conditioned *MES* resistance by the aphid effector MsEF1

By screening a Y2H library constructed from the mRNAs extracted from *MES* inoculated HN16 seedlings using RMES1A as a bait, a homolog of the PP5 phosphatases, which are conserved Ser/Thr-specific phosphatases in eukaryotes,^37,38^ was identified. This *MES* protein (MsEF1 hereafter), being composed of 493 residues and having a predicted molecular mass of 56.4 kDa, contained two tetratricopeptide (TPR) repeats and a putative calcineurin-like phosphoresterase domain (Figure 5A). MsEF1 homologs were present in diverse aphid species (Figure 5B). Although not studied in aphids, the fruit fly homolog of MsEF1, DmPP5, has been found to regulate oxidative stress mediated cell death.^39^

**Figure 5.**
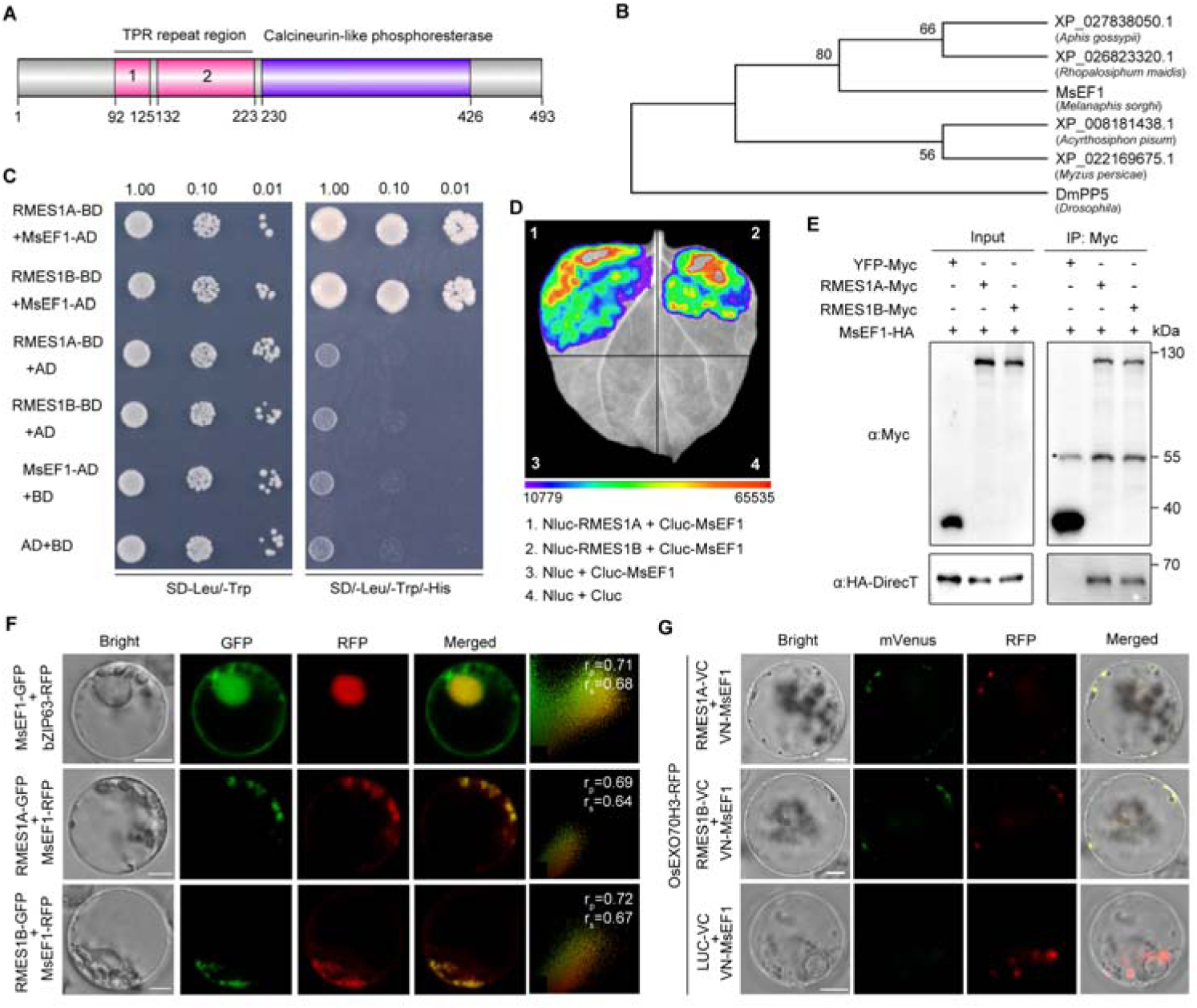
MsEF1 interacts with RMES1A and RMES1B in sorghum exocysts. (A) Schematic diagram of putative protein domains in MsEF1. (B) A neighbor-joining phylogenetic tree constructed using the amino acid sequences of MsEF1 and its homologs from diverse aphid species. (C) Y2H assays showing the interaction of MsEF1 with RMES1A and RMES1B. (D) Interaction of MsEF1 with RMES1A and RMES1B detected by SLC assays. The LUC signals were recorded at 48 h post infiltration of four different sets of expression constructs. (E) Co-IP assays verifying the interaction of MsEF1 with RMES1A and RMES1B in sorghum protoplasts. The indicated proteins were transiently expressed in protoplasts, which were then extracted and subjected to immunoblotting with anti-Myc and anti-HA-DirecT antibodies, respectively, for both the input and immunoprecipitated samples. YFP-Myc was used as a negative control. The asterisk indicates the heavy chain of IgG. (F) Subcellular localization of MsEF1 in sorghum protoplasts. bZIP63-RFP was expressed as a nucleus marker. Co-localization was evaluated by Pearson correlation coefficient (r_P_) and Spearman correlation coefficients (r_S_), where the value of +1.0 indicates complete colocalization. Scale bars, 5 µm. (G) MsEF1 interacts with RMES1A and RMES1B in sorghum exocysts as uncovered by BiFC assays. The three pairs of constructs, together with the one expressing OsEXO70H3-RFP, were introduced into protoplasts. Yellow fluorescence signals, resulting from reconstituted mVenus, signify MsEF1-RMES1A or MsEF1-RMES1B interaction. Red fluorescence signals caused by OsEXO70H3-RFP indicate the exocysts. Scale bars, 5 µm.

MsEF1 interacted with both RMES1A and RMES1B in yeast and tobacco cells (Figures 5C-5D). Such interactions were duly validated in sorghum protoplasts by Co-IP assays (Figure 5E). This led us to investigate which region(s) of RMES1A and RMES1B may be involved in interacting with MsEF1. RMES1A and RMES1B were each divided into N-terminal, middle, and C-terminal regions (Figure S6A). In Y2H and SLC assays, both the N-and C-terminal regions, but not the middle portion, interacted with MsEF1 (Figures S6B and S6C).

MsEF1 was present in both the cytoplasm and nucleus of sorghum protoplasts judging from the subcellular localization pattern of MsEF1-GFP fusion protein in the absence of RMES1A and RMES1B (Figure 5F). However, MsEF1 was relocated to sorghum exocysts according to the results obtained by co-expressing MsEF1-RFP with RMES1A-GFP or RMES1B-GFP (Figure 5F). BiFC assays showed the interaction of MsEF1 with RMES1A or RMES1B in the exocysts of sorghum protoplasts (Figure 5G), supporting relocation of MsEF1 to the exocysts in the presence of RMES1 proteins.

Importantly, we confirmed that RMES1A, RMES1B, and MsEF1 interacted with each other in sorghum cells by Co-IP assays (Figure 6A). Consistently, RMES1A, RMES1B, and MsEF1 interacted with each other in yeast tri-hybrid assays, with β-galactosidase activity level stimulated by three protein interaction (RMES1A-RMES1B-MsEF1) being significantly stronger than that by two protein interaction (RMES1A-RMES1B) (Figures 6B and 6C); co-expression of MsEF1 could also significantly boost the interaction between RMES1A and RMES1B in the SLC assays performed in tobacco leaves (Figure 6D). However, three protein interaction did not occur when RMES1A was replaced by the mutant asm1, neither could MsEF1 enhance the interaction between asm1 and RMES1B in yeast tri-hybrid and SLC assays (Figures 6B-6D). Collectively, these data suggest that MsEF1 is recruited to the exocysts in the presence of RMES1A and RMES1B, where it interacts with both RMES1A and RMES1B and strengthens their interaction. Mutation of RMES1A (e.g., asm1) disrupts the enhancement of RMES1A and RMES1B interaction by MsEF1.

**Figure 6.**
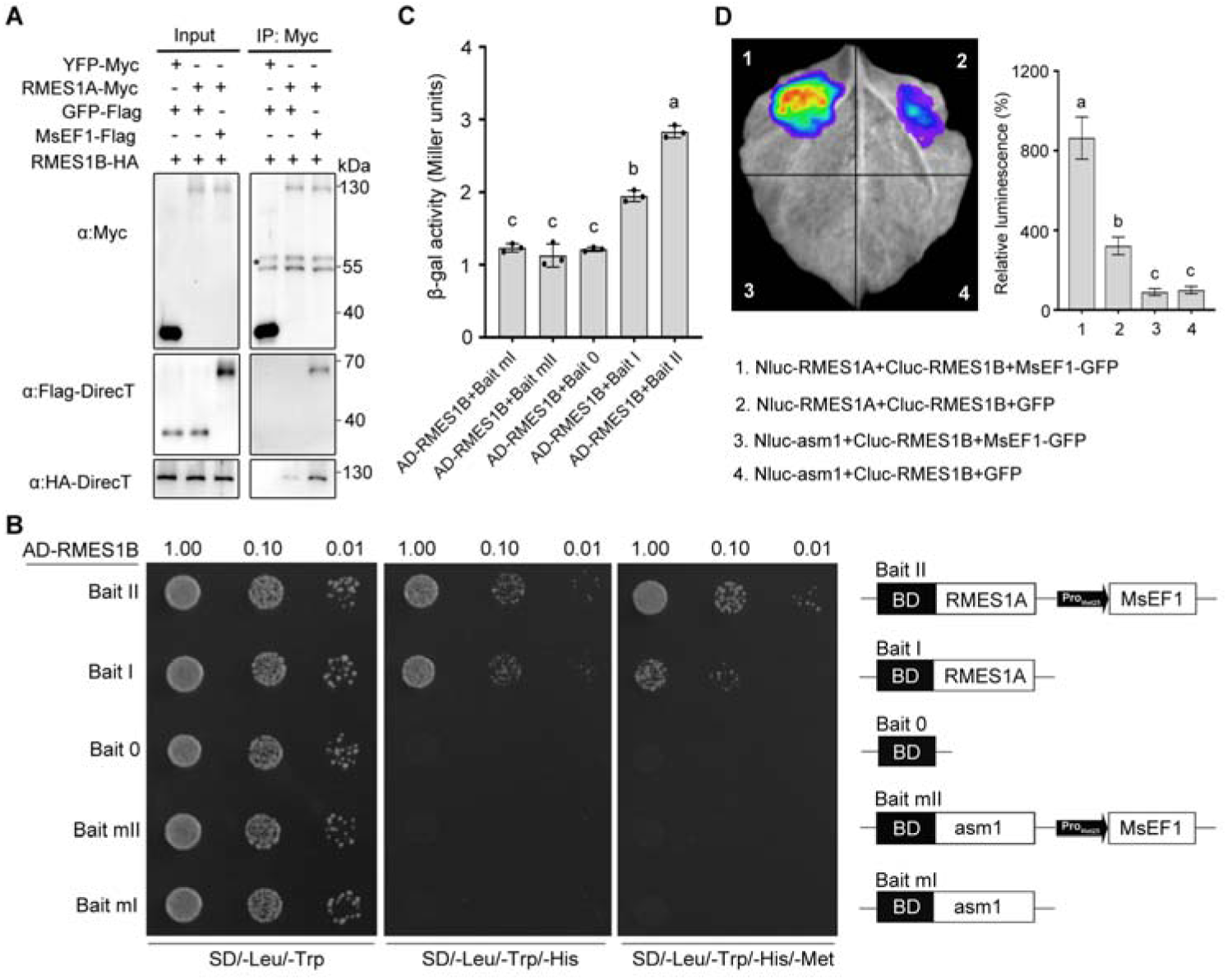
MsEF1 enhances the interaction of RMES1A and RMES1B. (A) MsEF1, RMES1A, and RMES1B form protein complex in sorghum as revealed by Co-IP assays. The proteins shown were transiently expressed in sorghum protoplasts, which were subsequently extracted and subjected to immunoblotting with anti-Myc, anti-HA-DirecT, and anti-Flag-DirecT antibodies, respectively, for both the input and immunoprecipitated samples. YFP-Myc served as a negative control. The asterisk marks IgG heavy chain. (B) Y3H assays showing that RMES1A-RMES1B interaction was enhanced in the presence of MsEF1. The bait and prey constructs used for Y3H assays were shown on the right panel. Note that the asm1 mutant failed to interact with RMES1B despite the presence of MsEF1. (C) Evaluation of interaction intensities for the different combinations of proteins examined in the Y3H assays in (B) using β-galactosidase assays. Error bars, means ± SE (n = 3). Dissimilar letters above the histograms indicate significant differences by Tukey’s HSD test (p < 0.05). (D) SLC assays validating the enhancement of RMES1A-RMES1B interaction by MsEF1 in tobacco foliar cells. The four sets of constructs used were listed in the bottom panel, with GFP expressed as a negative control (left panel). LUC signals were significantly stronger for set 1 constructs (+ MsEF1) than for set 2 constructs in which MsEF1 was replaced by GFP. No LUC signals were seen in the trials with RMES1A replaced by asm1 mutant. Relative intensities of LUC signals produced by the four sets of constructs were quantitatively compared (right panel), with those by set 4 constructs as control (100%). Error bars, means ± SE (n = 3). Dissimilar letters above the histograms indicate significant differences by Tukey’s HSD test (p < 0.05).

Finally, we used two approaches to investigate whether both RMES1A and RMES1B might be required for activating key anti-aphid defense response by MsEF1. First, epitope tagged RMES1A-HA, RMES1B-Myc, and MsEF1-Flag fusions were singularly, doubly, or triply expressed in the protoplasts of *MES* susceptible BTx623 followed by examining their effects on H2O2 production at 48 h post transfection. As shown in Figure S7A and 7B, H2O2 production was strongly enhanced in only the protoplasts with RMES1A-HA, RMES1B-Myc, and MsEF1-Flag simultaneously expressed, whereas no significant differences were detected in H2O2 content among the protoplast samples with GFP expression or with RMES1A-HA, RMES1B-Myc, and MsEF1-Flag singularly or doubly expressed. Second, we purified a bacterially expressed, MBP-tagged MsEF1 fusion protein (MBP-MsEF1, Figure S7C), and pressure-injected it into the leaves of HN16, NIL^R+^, five *asm* mutants, and BTx623, respectively. Based on DAB staining and quantitative determination of H2O2 at 48 hours post injection, stronger H2O2 burst (relative to that observed for MBP injection) was found in the leaves of HN16 and NIL^R+^ but not those of BTx623 and the five *asm* mutants (Figure S7D and S7E). These lines of evidence support the requirement of both RMES1A and RMES1B for MsEF1 to promote ROS production, an important anti-aphid defense response.

### A new type of resistance proteins represented by RMES1A and RMES1B

RMES1A and RMES1B displayed low amino acid sequence identities (< 30%) to either CNLs or TNLs, but their identities to BPH6 and BPH30, two previously reported LRR-containing proteins conferring BPH resistance in rice,^12,35^ were above 30% (Table S2). Consistently, phylogenetic analysis showed that RMES1A, RMES1B, BPH6, and BPH30 formed a cluster separated from those composed of typical CNLs or TNLs (Figure S8). No homologous CC, TIR, NB domains were detected in RMES1A, RMES1B, BPH6, and BPH30 (Figure S8), when their amino acid sequences were compared to those of two representative structurally-resolved NLRs, i.e., SR35 (CNL), and RPP1 (TNL), which controlled the immunity to plant pathogens.^40,41^

Hence, we performed protein structure modeling using the AlphaFold2 system.^42^ RMES1A was predicted to have four domains with a relatively high predicted local distance difference test (pLDDT) score (Figures 7A-7C). The N-terminal domain (M1-P194) was composed of mainly α-helices and β-strands, the middle domain (F195-Y299) contained largely α-helices, and the C-terminal region (T300-R1032) carried two LRR domains, which were separated by three pairs of antiparallel β-strands and had their concave sides opposing to each other (Figures 7A and 7B). The presence of two LRR domains in BPH30 was also suggested by Shi et al. (2021) based on protein annotation information in the InterPro database. The two predicted LRR domains in RMES1A may form a wave-shaped structure, which differs from the single, horseshoe-shaped LRR domain present in SR35 and RPP1 (Figures 7B). Similar prediction results were obtained for the N-terminal and middle domains and the C-terminal region of RMES1B, BPH6, and BPH30 (Figures 7C and S9A).

**Figure 7.**
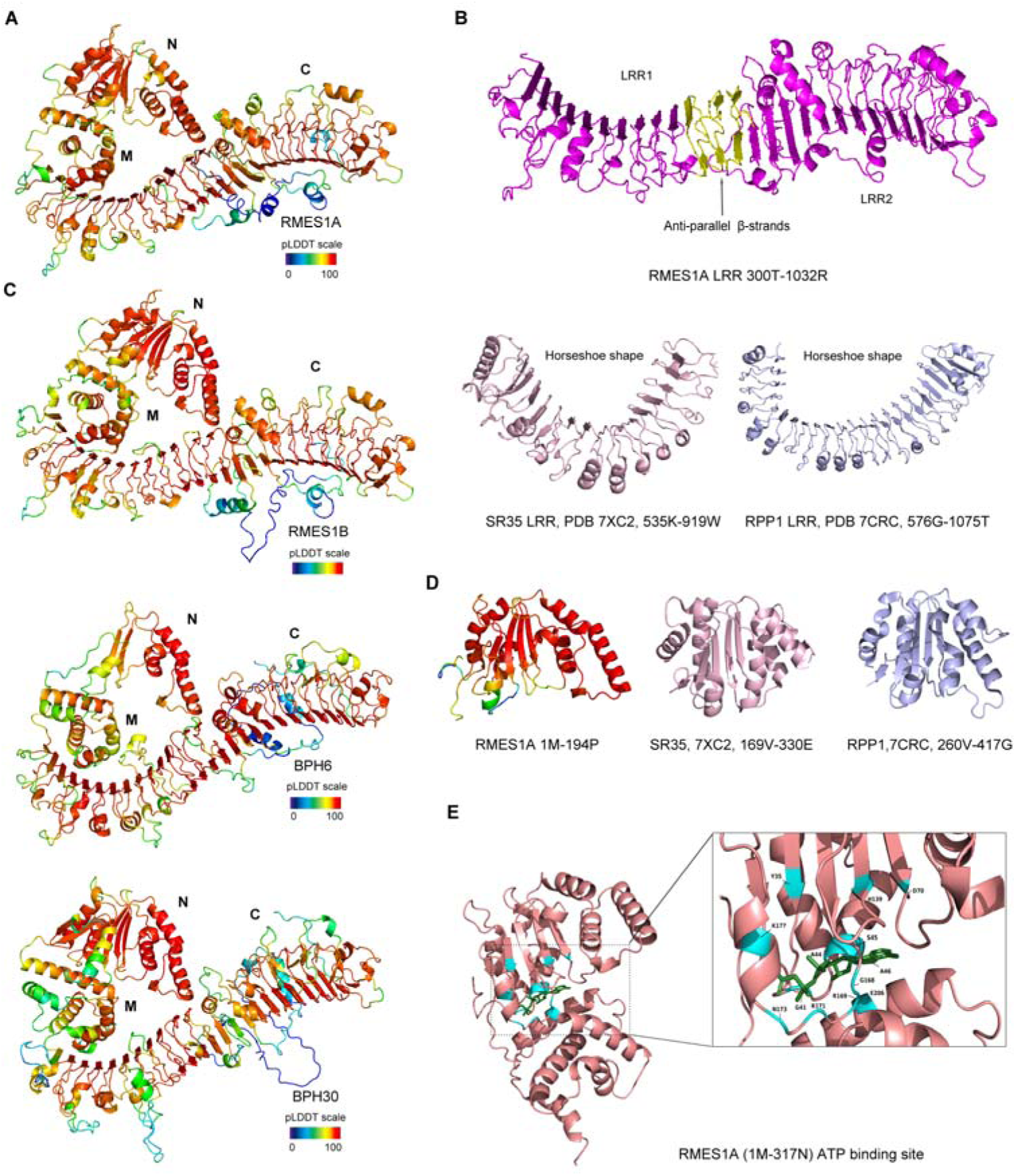
Structural modeling of RMES1A, RMES1B, BPH6, and BPH30 using the AlphaFold2 system. (A, B) The overall structure of RMES1A predicted by AlphaFold2 (A). Four putative domains could be discerned, i.e., N-terminal domain (N), middle domain (M), and two LRR domains in the C-terminus (C) (A, B). The C-terminus of RMES1A, with two probable LRR domains having opposing concave sides and separated by three pairs of anti-parallel β-strands, forms a wave-like structure, which differs from the horseshoe-shaped LRR domain of SR35 and RPP1. The pLDDT scale shows prediction accuracy, with 100 indicating high prediction quality. (C) The overall predicted structure of RMES1B, BPH6, and BPH30 bears high similarity to that of RMES1A. (D) The putative N-terminal domain of RMES1A showing similarity with the nucleotide binding domain of SR35 and RPP1. (E) Prediction of a potential ATP binding site in a grove between the N-terminal and M domains of RMES1A. The residues possibly involved in ATP binding are labelled.

The putative N-terminal domain of RMES1A resembled the NB domain of SR35 and RPP1, with the RMSD value reaching 6.027 and 4.899, respectively (Figure 7D). Interestingly, an ATP binding site was predicted in a groove between the N-terminal and middle domains, with 12 residues (Y35, G41, A44, S45, A46, D70, H139, G168, R169, R171 and K177 in the N-terminal domain; E206 in the middle domain) potentially involved in ATP binding (Figure 7E). The predicted N-terminal domain in RMES1B, BPH6, and BPH30 also resembled the NB domain in SR35 and RPP1 (Figure S9B), and ATP binding site was predicted to exist in RMES1B, BPH6, and BPH30 as well (Figure S9C). No CC or TIR domain was predicted for RMES1A, RMES1B, BPH6, and BPH30, in line with the result presented in Figure S8. Finally, the asm1 mutant missing 59 residues in the C-terminal region lacked two LRR repeats in the second predicted LRR domain of RMES1A (Figure S9D). Together, these data suggest that RMES1A, RMES1B, BPH6, and BPH30 represent a new type of resistance proteins carrying a potential NB domain in the N-terminus and two putative LRR domains in the C-terminus.

## DISCUSSION

There is currently an urgent need to develop *MES* resistant sorghum cultivars to limit the mounting damages caused by rapid expansion of *MES*.^19–22^ *RMES1*, initially identified and mapped by us over 10 years ago,^31,43^ has been verified to confer *MES* resistance in many research and breeding studies.^19,22,29,30,44–47^ To facilitate more effective and rational use of *RMES1*, we cloned the causal genes of *RMES1* and studied the genetic and molecular mechanisms underlying their function.

By integrating the different findings made in this work, the main functional characteristics of *RMES1* and the underlying mechanisms can be summarized as follows. In resistant cultivar, *MES* feeding enhances the expression of *RMES1A* and *RMES1B* in leaf sclerenchyma and vascular cells, resulting in elevated protein abundance of RMES1A and RMES1B in the exocysts. The aphid effector MsEF1 is recruited to the exocyst complex where it binds to RMES1A and RMES1B and strengthens their interaction. This activates immune signaling, leading to upregulation of defense responses including ROS burst and callose deposition. Eventually, these processes give rise to both antixenosis and antibiosis resistance, which not only deters *MES* settling on the resistant plant but also inhibits its reproduction. In the *ams1* mutant, disruption of the second LRR domain renders *MES* susceptibility via abrogating the interactions among RMES1A RMES1B, and MsEF1 (Figures 1, 4, 6, S1 and S9). Tetreault et al. (2019) showed the operation of both antixenosis and antibiosis in the *MES* resistant sorghum line RTx2783 that carries *RMES1* locus.^29^ VanGessel et al. (2023) found that *MES* resistance conditioned by *RMES1* in the sorghum line IRAT204 involved stronger antibiosis but weaker antixenosis.^22^ These data provide direct support to our finding obtained using *RMES1A* and *RMES1B* mutants (Figure S4).

Compared to previous studies on insect resistance genes, our work is unique in revealing a pair of tandemly linked genes both required for insect resistance. This is reminiscent of the findings of many paired NLRs controlling plant resistance to pathogenic microbes. In such systems, the sensor and helper NLRs interact with each other, with the former recognizing pathogen avirulent effector and the latter initiating immune signaling.^6,7^ A MADA motif, MADAxVSFxVxKLxxLLxxEx (with x indicating non-conserved amino acid residues), has been detected in a subset of helper CNLs but not in their sensor CNLs.^48^ We examined the amino acid sequences of RMES1A and RMES1B and found no MADA motif in either of them. This is in line with our structural modelling result that did not find the presence of CC or TIR domain in RMES1A and RMES1B (Figures 7 and S9). Hence, the activation and signaling mechanisms of RMES1A/RMES1B pair differ from those of previously studied CNL and TNL pairs. Discovery of RMES1A and RMES1B as a functional pair expands the spectrum of molecularly characterized insect resistance proteins, whose continued study will contribute new knowledge to the mechanisms behind the functions of paired resistance proteins in plants.

We found that MsEF1 interacted with both the putative N-terminal ATP binding domain and the C-terminal LRR region (Figure S6). This contrasts with typical plant pathogen-derived avirulent effectors that generally bind to only the LRR domain (SR35) or the LRR and C-JID domains (RPP1 and ROQ1) of NLRs,^40,41,49^ and indicates potential novelty in the activation of RMES1A and RMES1B by MsEF1. Furthermore, MsEF1 is a homolog of the PP5 phosphatase that is conserved in eukaryotic cells and regulates diverse cell growth and death processes under normal or stress conditions,^37,38^ but to our knowledge no microbial or insect PP5 proteins have previously been shown to act as avirulent or virulent effectors. Considering that PP5 proteins have been shown to possess both protein phosphatase and molecular chaperone activities,^37,38^ it will be interesting to investigate whether the activation of RMES1A and RMES1B by MsEF1 may involve dephosphorylation and/or protein folding events in further research. The identification of MsEF1 in this work broadens the functional understanding of eukaryotic PP5 phosphatases and the types of effectors activating plant resistance proteins, as PP5s have not been reported to act as resistance protein activation effectors before.

The exocyst, a conserved multiprotein complex in eukaryotic cells,^50,51^ is emerging as a critical battle field for attack and counteract warfare in plant biotic interactions.^52^ Many pathogens have evolved virulent effectors to suppress the function of exocyst complex by interacting with its constituent subunits, e.g., Exo70s, which compromises the delivery of defense related components to the plasma membrane or extracellular space and thereby leads to disease susceptibility.^53–56^ In turn, plants have developed NLR based resistance that is initiated after detecting structural and/or functional alterations of exocysts caused by pathogen effectors.^57^ Here we demonstrate that RMES1A, RMES1B and their avirulent effector MsEF1 are all localized to the exocyst complex where they interact with each other to trigger immune signaling, thus providing an efficient system for further studying the role of exocysts in plant immunity against insect pests. MsEF1 was distributed in both the cytoplasm and nucleus in the absence of RMES1A and RMES1B but were relocated to the exocyst upon the presence of the two proteins (Figure 5F). This suggests that RMES1A and RMES1B may directly recruit MsEF1 to the exocyst through protein-protein interaction. Alternatively, certain exocyst subunit(s) may interact with RMES1A and RMES1B as well as MsEF1, thus making it possible for the three proteins to accumulate and to interact with each other in the exocyst complex to activate immune signaling.

Recently, the discoveries of CNL resistosome acting as calcium permeable channel and TNL resistosome as NADase to generate nucleotide based second messengers have provided novel directions for further dissecting the molecular and biochemical basis of plant disease resistance.^40,41,49,58–61^ Here we predicted the secondary structures of RMES1A, RMES1B and their homologs BPH6 and BPH30 with relatively high confidence (Figures 7 and S9). Presence of potential ATP binding site and two LRR domains, but lacking typical CC and TIR modules, in their predicted structures indicates that the four proteins likely represent a new type of atypical NLRs in plants. Cryo-electron microcopy is underway to elucidate the protein complex formed by RMES1A, RMES1B, and MsEF1, which may yield new knowledge on the structure and activation mechanism of plant resistance proteins. We note that homologs of the four proteins are widely present in monocot plants, most of which are annotated as putative resistance proteins (Table S3). The insights generated in this work, together with those reported for BPH6 and BPH30,^12,35^ will stimulate more systematic studies on this new type of resistance proteins.

Finally, the characterization of RMES1A and RMES1B and their avirulent effector MsEF1 will enhance efficient breeding and effective use of *MES* resistant sorghum cultivars in several ways. First, the PCR primers used for amplifying *RMES1A* or *RMES1B* coding sequence can be converted into DNA markers to speed up the transfer of both genes into desirable genetic backgrounds. Second, sorghum cultivars carrying *RMES1* can be planted in the regions dominated by MsEF1 expressing *MES* population. In other regions where the *MES* population does not express MsEF1 or express the MsEF1 with mutations, *RMES1* carrying sorghum cultivars should be avoided. These practices can extend the durability of *RMES1* conditioned resistance and avoid severe sorghum crop losses due to sudden resistance breakdown. Third, the 36 *MES* resistant accessions with functional *RMES1A* and *RMES1B* genes, originated from China, India, Japan, and USA (Table S1), can contribute to breeding of *MES* resistant sorghum by serving as parental materials. Finally, our characterization of *RMES1A* and *RMES1B* is useful for clarifying functional relationship between *RMES1* and *RMES2*. The knowledge gained will facilitate appropriate pyramiding of the two loci for sustainable control of *MES* infestations in global sorghum production.

## ACKNOWLEDGMENTS

We thank Professors Haichun Jin, Hongwei Cai, and Zhiying Ma for constructive suggestions on this work, Dr. Zhengqing Fu for critical reading of manuscript, and staff at the Analytical Facility Center of Henan Agricultural University for assistance on confocal microscopy. This work was supported by the National Natural Science Foundation of China (32301770), the Ministry of Agriculture of China (2016ZX08009003-001), Henan Provincial government (SN01-2022-01), and Hebei Province Key Research & Development Program (21326305D).

## AUTHOR CONTRIBUTIONS

D.-W.W., K.Z., B.Y., K.L., and D.W. conceived and designed the experiments. K.L., D.T., Y.S., F.W., J.C., D.W., Z.D., J.-C.C, S.Z., S.N.C., J.J.L., H.L., and M.W. performed the experiments and analyzed the data. G.-Q. L., and P.L. collected and maintained the 166 sorghum accessions. G.L., K.L., and D.W. conducted bioinformatic analysis. J. Z. and X.J. carried out protein structure modeling. D.-W.W., Q.S., B.Y., D.W., and K.L. wrote the manuscript. All authors discussed and contributed to the manuscript.

## MATERIALS AND METHODS

### Plant materials and growth conditions

The sorghum materials used included HN16, BTx623, five *asm* mutants, two NILs (NIL^R+^ and NIL^R-^), P89002 (PI 561847), TX430 (PI 655996), and 166 sorghum germplasm accessions. For field planting, sorghum genotypes (populations) were sown in May and harvested in October using standard cultivation practices. Alternatively, sorghum plants were cultivated in a greenhouse set at 28 ℃ with 60% relative humidity and a 16-h-light/8-h-dark photoperiod. For investigating the response of different sorghum materials to *MES* infestation at the seedling stage, the desired genotypes were grown in growth chamber with environmental parameters set as above. Tobacco (*Nicotiana benthamiana*) plants were raised in the greenhouse at 25 ℃ with 60% relative humidity and a 16-h-light/8-h-dark photoperiod.

### Insects

Sorghum aphid, previously named as *Melanaphis sacchari* but now corrected as *Melanaphis sorghi*^20,21^, was reared on the seedings of BTx623 at 28 ℃ under a 16-h-light/8-h-dark photoperiod with 60% relative humidity.

### Fine mapping of *RMES1*

Two flanking markers Sb6m2650 and Sb6rj2776,^31^ were used to screen 5,136 BC_5_F_2_ plants, resulting in 44 recombinants. Additional DNA markers were designed based on BTx623 genome sequence, which were used to analyze the recombinants. This narrowed *RMES1* to a 25 kb region between markers Sb6id2705 and Sb6id2730. A bacterial artificial chromosome (BAC) library was constructed for the aphid resistant sorghum cultivar HN16 as described previously.^62,63^ The library, containing 57,600 clones and covering approximately 6.25× sorghum genome, was screened by PCR as previously detailed.^64^ Ten positive BAC clones were identified. Four minimum tilling path BAC clones covering *RMES1* region were sequenced by MiSeq and their inserts were assembled. A 194 kb sequence was obtained, with four LRR protein encoding genes, i.e., *LRRP2*, *LRRP3*, *RMES1A* and *RMES1B*, found on a subregion of 169 kb, which corresponded to the 25 kb interval bordered by Sb6id2705 and Sb6id2730.

### Development of *asm* mutants and NILs

Diepoxybutane was used to mutate 3,000 mature seeds of HN16 as described previously,^65^ with 1,500 M_2_ families cultivated in the field and screened for *MES* susceptible mutants by visual inspection of *MES* aphids on the leaves. Aphid susceptible mutant (*asm*) was found in eight different families from which *asm1* was identified by genomic PCR amplification of *RMES1A* or *RMES1B* coding sequence. Subsequently, ethyl methane sulphonate was used to treat 10,000 HN16 seeds,^65^ with 3,176 M_2_ families grown in the field and checked for *asm* mutants by visual inspection as above. The resultant aphid susceptible M_2_ plants were investigated for potential mutation in *RMES1A* or *RMES1B* coding sequence by genomic PCR assays, leading to the identification of *asm2*. Besides, 10,000 HN16 seeds were irradiated with two doses of fast neutron (10 and 20 Gy, 5,000 seeds each),^66^ with 7,800 M_1_ families obtained. These families were screened in the greenhouse by inoculating *MES*, which allowed the identification of *asm3* and *asm4* in two independent families. The two mutants were found to lack *RMES1B* by genomic PCR amplifications as above. To clarify the precise chromosomal deletions occurred in *RMES1* locus in *asm3* and *asm4*, the two mutants were resequenced in HiSeq 2,000 platform,^67^ with 26.81 Gb and 25.06 Gb of clean data obtained for *asm3* and *asm4*, respectively. After removing low quality reads and adaptor sequences, the reads were mapped to the 169 kb HN16 BAC contig covering *RMES1* locus. Read depth was calculated with mosdepth (v.0.2.6) using 100 bp non-overlapping sliding windows. This revealed the low coverage region that represented the deleted segment containing *RMES1B*. Lastly, the double mutant *asm1asm4* was prepared by artificial crossing between *asm1* and *asm4*.

A BC_7_F_1_ plant was selfed in the greenhouse. The resultant BC_7_F_2_ population was screened for the plants with or without *RMES1* locus by amplifying the coding sequence of *RMES1A* and *RMES1B*, leading to the development of NIL^R+^ and NIL^R-^, respectively.

### Generation of CRISPR mutant of *RMES1A* and *RMES1B*

Genome editing of *MES* resistant sorghum was performed using the sorghum line P89002, which was confirmed to carry *RMES1* locus by PCR amplification and DNA sequencing of the coding region for *RMES1A* and *RMES1B*, respectively. The CRISPR/Cas9 method was used for the editing.^68^ The sgRNA target sites were 5′-GCACTGATATGGTTGACT-3′, and 5′-GAAATGGGTGGCAGTACCA-3′, which were specifically conserved among *RMES1A* and *RMES1B*. Five sibling lines, derived from a homozygous mutant (P89002 #25) with knockout mutations in both *RMES1A* and *RMES1B*, were tested for the response to *MES* infestation. The mutations in *RMES1A* and *RMES1B* in P89002 #25 and its derivative lines were validated by Sanger sequencing of the relevant PCR amplicons with primers specific for *RMES1A* and *RMES1B*, respectively (Table S4).

### Assessing the responses of *asm* and CRISPR mutants, *RMES1* NILs, and 166 germplasm accessions to *MES*

The seeds were germinated in the growth chamber at 28 ℃ (see above). At two-leaf stage, 15 well grown seedlings (5 in each pot) per genotype were inoculated with apterous adult aphids (10 per seedling unless otherwise stated). After inoculation, each pot was covered with an insect proof device, followed by returning to the growth chamber. The mean number of aphids for each genotype was determined at 24, 48, 72, 96, 120, and/or 168 h post inoculation using the values collected from 15 seedlings (in three separate experiments). In general, the susceptible genotypes (e.g., BTx623, TX430, *asm* mutants, CRISPR knockouts, and NIL^R-^) became dying at 2 weeks after inoculation, while the resistant materials (e.g., HN16, P89002, and NIL^R+^) continued to grow.

### Free choice and no-choice tests and honeydew assays

These experiments were conducted with sorghum seedlings at two-leaf stage in the growth chamber at 28 ℃ and using adult apterous *MES* aphids. In a free choice test, 8 trials were conducted, with each involving two desired genotypes (one resistant and one susceptible, 4 seedlings each and cultivated in the same pot). Each seedling was inoculated with 5 aphids, with the number of aphids remained on each seedling recorded at 3, 6, 24, 48 and 72 h after inoculation. In a no-choice test, 8 sets of trials were carried out, with each involving one resistant and one susceptible genotype. For each genotype, 8 seedlings were used, with 10 aphids inoculate on each seedling. The number of nymphs produced by the aphids on each seedling was at scored at 3, 6, 24, 48 and 72 h after inoculation.

The honeydew secretion assay was conducted as previously described.^69^ Two fresh stem segments (1.5 cm each) of the desired sorghum genotype were placed onto the filter paper in a clean petri dish, with 5 aphids inoculated and lid covered. The whole device was transferred into a growth chamber set at 28 ℃ with 80% humidity. The aphids and stems were discarded at 36 h post inoculation, with the filter paper stained using 0.1% (w/v) ninhydrin in acetone solution. The area of ninhydrin sediment was measured using the IMAGEJ software.

### RT-qPCR assays

Total RNAs were extracted using the RNAiso Plus reagent (TaKaRa, Ostu, Japan), which were converted into cDNA with Superscript II reverse transcriptase (Invitrogen, Carlsbad, CA) according to manufacturer’s instruction. The cDNA was diluted and used as templates for quantitative PCR with gene-specific primers in a Light Cycler 480 Real Time PCR system (Roche, Bassel, Switzerland). Gene-specific primers were listed in Table S4. All RT-qPCR assays were repeated three times independently. The amplification of a sorghum actin gene (NCBI accession: XM_002441128) served as an internal control for all RT-qPCR assays.

The expression profiles of *RMES1A* and *RMES1B* during sorghum development was investigated using RT-qPCR assays with total RNA samples extracted from the roots, stems, leaves, flag leaves, spikes, and mature grains of greenhouse cultivated HN16 plants at the heading or harvesting stage. For investigating the expression changes of *RMES1A* and *RMES1B* in response to *MES* feeding, HN16 seedlings (at two leaf stage) were inoculated with *MES* (20 adult aphids per plant), with their leaf sheaths collected at 6, 12, 24, 48, and 72 h after *MES* inoculation, respectively. They were then used in RT-qPCR assays as stated above.

### *In situ* hybridization assays

HN16 seedlings were inoculated with *MES* as above. Their leaf sheaths, collected at 48 h after the inoculation, were cut into 1 cm segments, followed by fixation in FAA solution (3.7% form aldehyde, 5% acetic acid, 50% ethanol). Leaf sheaths were also collected from uninoculated HN16 seedlings, which were similarly fixed as controls. The fixed samples were processed for wax embedding and sectioning for *in situ* hybridization assays as reported in previous studies.^70^ The antisense probe used in the assays was prepared using the oligonucleotide 5′-UGCUUCUACGGCUUCAUUGAUUCU-3′, which was specific for the coding sequence of *RMES1A* and *RMES1B*. As a control, the complementary sequence of this oligonucleotide was employed to prepare a sense probe. All probes were labeled with digoxigenin-dUTP, with hybridization signals detected using DAB as a substrate.^71^ The hybridized sections were examined under a light microscope equipped with a digital camera (Zeiss, Jena, Germany).

### Y2H and Y3H assays

The GAL4 yeast two-hybrid system was used to investigate protein-protein interactions. The coding sequence of *RMES1A*, *RMES1B* or derivative fragments was each amplified and cloned at *EcoR*I and *BamH*I sites of pGBKT7 (BD) or pGADT7 (AD) vectors as desired. The coding sequence of MsEF1 was cloned at *EcoR*I and *BamH*I of pGADT7 (AD). Different pairs of the constructed vectors were co-transformed into the yeast strain Y2Hgold and grown on synthetic medium lacking leucine and tryptophan (SD/-Leu/-Trp), with the resultant clones verified for protein-protein interaction on the medium lacking leucine, tryptophan, and histidine (SD/-Leu/-Trp/-His) at 30 ℃ for 3 d. For Y3H assays, the coding sequence of RMES1A or asm1 was specifically amplified and cloned into the MSC I location of the pBridge vector at the *EcoR*I-*BamH*I sites, resulting in Bait constructs. The coding sequence of MsEF1 was amplified and cloned into the MCS II location of Bait I/Bait mI (pBridge-RMES1/pBridge-asm1) at the *Not*I sites, producing Bait II/Bait mII. The construct pGADT7-RMES1B, developed in Y2H assays, was used as prey. Various pairs of constructs were co-introduced into the yeast strain AH109, with the cells grown on SD/-Leu/-Trp medium for 3 to 5 d. The yeast strains, carrying different sets of constructs, were serially diluted, followed by plating onto the SD/-Leu/-Trp/-His and SD/-Leu/-Trp/-His/-Met media, respectively.

The O-nitrophenyl-β-d-galactosidase (ONPG) tests in Y3H assays were performed as described by Kippert (1995).^72^ Briefly, yeast cells were resuspended with 800 μl of Z-buffer (60 mM Na_2_HPO_4_, 40 mM NaH_2_PO_4_, 10 mM KCl, 1 mM MgSO_4_, and 50 mM β-mercaptoethanol) and placed on ice. Then the suspension was incubated at 30 ℃ for 15 min, followed by the addition of 160 µL of 4 mg mL^−1^ O-nitrophenyl-β-d-galactoside. The mixture was briefly shaken and incubated at 30 ℃ for 15 min. Afterwards, the reaction was stopped by adding 400 μL of 1 M Na_2_CO_3_. After centrifugation for 30 s at 16,000 g, the OD_420_ value of the supernatant was determined in a spectrophotometer (Thermo Scientific, Waltham, USA).

### Identification of MsEF1

MsEF1 was identified by Y2H screening of a cDNA library prepared using the mRNAs derived from approximately 6,000 adult *MES* aphids that had been feeding on HN16 seedlings for 2 days. The Y2H library, containing 2 × 10^6^ recombinant clones, was developed using the SMART cDNA Library Construction Kit (Clontech, Code No.634901) as described by He et al., (2019).^73^ A bait vector pGBKT7-RMES1A, with the full-length coding sequence of RMES1A fused to that of DNA binding domain, was used to screen the Y2H library. Co-transformed yeast cells were grown on the SD/-Leu/-Trp/-His medium for 3-5 days at 30 ℃. The resultant positive clone carrying MsEF1 coding sequence was further analyzed and validated for interactions with RMES1A and RMES1B using additional methods including the SLC, BiFC, and Co-IP assays conducted in tobacco foliar cells or sorghum protoplasts (see below).

### Immunoblotting and Co-IP assays

Total proteins were extracted from the sorghum protoplasts transfected with desired expression constructs using the extraction buffer (100 mM Tris-HCl, pH 7.5, 5 mM MgCl_2_, 1 mM EDTA, 0.5% Triton X-100, 1 mM DTT, and 1 mM PMSF). The resulting proteins were separated using 10% SDS-PAGE for immunoblotting as detailed in Jin et al. (2022).^74^ For detecting the target proteins fused with Flag, HA, or Myc tag, appropriate commercial antibodies were used in 1:2,000 dilution. The second antibody was goat anti-mouse IgG conjugated to horseradish peroxidase, which was used in 1:5,000 dilution. The immunosignals were recorded in a chemiluminescence detector (Tanon 5200, Shanghai, China).

For Co-IP assays, the coding sequence of RMES1A, asm1, RMES1B, or MsEF1 was each cloned into the vector pHZ205-Myc, pHZ204-HA, or pHZ206-Flag to generate the ubiquitin gene promoter driven cassettes for expressing RMES1A-Myc, asm1-Myc, RMES1B-HA, or MsEF1-Flag in the protoplasts of BTx623. Protoplast preparation and transfection were executed as reported by Zhang et al. (2011).^75^ The transfected protoplasts were incubated for 24 h at 28 ℃ in the dark, followed by centrifugation at 450 g at 4 ℃ for 5 min. Total proteins were extracted from the pelleted protoplasts with 300 µl extraction buffer as described above. Immunoprecipitation was accomplished by adding 20 µl anti-Myc antibody coupled protein G agarose, with the mixture incubated for 5 h at 4 °C. The immunoprecipitated proteins were separated using 10% SDS-PAGE, which were subjected to immunoblotting assays using anti-Myc, anti-Flag, anti-HA, or anti-HA-HRP-DirecT antibody to check the expression of target proteins in input samples or the presence of interacting protein(s) in the immunoprecipitates.

### Subcellular localizations

The coding sequence of target genes (*RMES1A*, *RMES1B*, and *MsEF1*) was amplified using gene-specific primers and cloned into the pRHVcXFP (GFP or RFP) vectors carrying ubiquitin gene promoter directed expression cassette.^76^ The resulting constructs were introduced into BTx623 protoplasts as indicated above, with the expressed GFP or RFP fusion proteins examined at 24 h after transfection under a confocal microscope (Zeiss LSM710, Jena, Germany). The C-terminal RFP fusion of bZIP63,^77^ or that of OsEXO70H3,^36^ was also expressed as a nucleus or exocyst marker. For assessing the degrees of co-localization, Pearson and Spearman correlation coefficients were calculated by analyzing 30 individual images using the IMAGEJ software supplemented with the PSC colocalization program.^78^ GFP fluorescence was examined using excitation at 488 nm (emission at 507-550 nm), while RFP signals were visualized with excitation at 561 nm (emission at 588-630 nm).

### BiFC assays

The pRHV suit of BiFC vectors, developed by He et al. (2018),^76^ were used to generate the constructs expressing VN-MsEF1, VN-asm1, VN-RMES1A, RMES1A-VC, and RMES1B-VC, respectively. The desired constructs were introduced into BTx623 protoplasts along with that expressing OsEXO70H3-RFP. The protoplasts were examined by confocal microscopy at 24 h post transfection as detailed above, with the excitation wavelength being 514 nm (for mVenus) or 561 nm (for RFP).

### SLC assays

SLC assays were performed in tobacco leaves as described previously.^79^ Briefly, the coding sequence of *RMES1A*, *RMES1B*, or *MsEF1* was cloned into pCAMBIA1300-NLuc or pCAMBIA1300-CLuc vector. The resulting NLuc and CLuc constructs were each introduced into the *Agrobacterium tumefaciens* strain GV3101. The recombinant strains were introduced into the leaves of four-week-old tobacco in desired combinations through agroinfiltration, with luciferase signals detected at 48 h after the infiltration using a NightShade LB985 Plant Imaging System installed with the IndiGO software for recording and quantifying LUC signals (Berthold, Bad Wildbad, Germany).

### DAB staining and measurement of H_2_O_2_

DAB staining of sorghum leaf samples was executed as outlined by Daudi and O’Brien (2012).^33^ H_2_O_2_ content was measured as reported previously using a commercial kit (Molecular Probes, OR, USA).^70^

### Callose staining and quantification

Callose deposition induced by *MES* feeding was examined by staining with 0.01% aniline blue, with the staining signals recorded using confocal microscopy.^80^ Callose contents of the sorghum leaf samples collected before or after *MES* feeding were determined using a callose assay kit (Keming, Suzhou, China).

### Purification and pressure injection into sorghum leaves of MBP-MsEF1

*MsEF1* coding sequence was cloned into the vector pMAL-c2X, which was expressed in *E. coli* and purified as a MBP (maltose binding protein)-fusion protein.^81^ MBP was similarly expressed and purified as a control. The purified MBP-MsEF1 and MBP proteins were diluted to 0.2 mg/ml in 1× PBS for pressure injection into the leaves of different sorghum genotypes (Figure S7D). In this set of assays, 25 days old sorghum plants raised in the growth chamber were used, with MBP-MsEF1 and MBP proteins (50 μl each) pressure-injected into the opposite sides of the main vein as detailed in Deng et al. (2022).^81^ The infiltrated areas were excised at 36 h post injection for determining H_2_O_2_ by DAB staining or quantitative measurement as outlined above.

### Bioinformatic and phylogenetic analyses

*MsEF1* homologs in different aphid genomes were searched in the NCBI database (https://www.ncbi.nlm.nih.gov/) using MsEF1 sequence as query. *RMES1A* and *RMES1B* homologs were searched similarly using RMES1A and RMES1B sequences as queries. The amino acid sequences of representative CNLs and TNLs and previously cloned insect resistance genes were also retrieved from NCBI. Their GenBank accessions are AEE78731.1 (ZAR1), AEE77906.1 (RPP1), ATD14363.1 (ROQ1), OAO91745.1 (RPS4), AGP75918 (SR35), AAC67238.1 (Mi-1), AIU36098 (Vat), ASV63918.1 (BPH6), ANC90314.1 (BPH9), ADB07392 (BPH14), and QYA53811.1 (BPH30). Sequence identity and similarity were computed using MatGAT.^82^ MsEF1 protein domain was predicted using HMMER (https://www.ebi.ac.uk/Tools/hmmer/search/hmmscan). All amino acid sequence alignments were conducted using ClustalW in the MEGA 11 software using neighbor-joining method.^83^

### Protein structural modeling

The full-length proteins of RMES1A, RMES1B, BPH6, and BPH30 were structurally modelled using AlphaFold 2.0 (https://colab.research.google.com/github/sokrypton/ColabFold/blob/ main/AlphaFold2.ipynb#scrollTo=kOblAo-xetgx).^42^ ATP binding site prediction was performed with the molecular docking program DiffDock at https://github.com/gcorso/diffdock,^84^ followed by analysis using Protein-Ligand Interaction Profiler (PLIP) at https://plip-tool.biotec.tu-dresden.de/plip-web/plip/index.^85^ Comparison and analysis of protein structures were accomplished with the aid of Pymol V2.2.0 (https://pymol.org/2/).

### Statistical analysis

All assays were repeated three times using independent biological replicates, with each replicate having at least three technical repeats. Numerical values were presented as means ± SE. Statistical analysis of the data was performed using Student’s *t* test or Tukey’s HSD test in the Prism graphic software (Graph Pad Software, version 8.1.2, San Diego, CA, USA).

## SUPPLEMENTAL INFORMATION

Supplemental Figures with legends. Figures S1–S9 with legends.

Supplemental Tables. Tables S1-S4.

