## Supplemental figures for "Activation of two noncanonical R proteins by an insect effector confers plant immunity to aphid infestation"

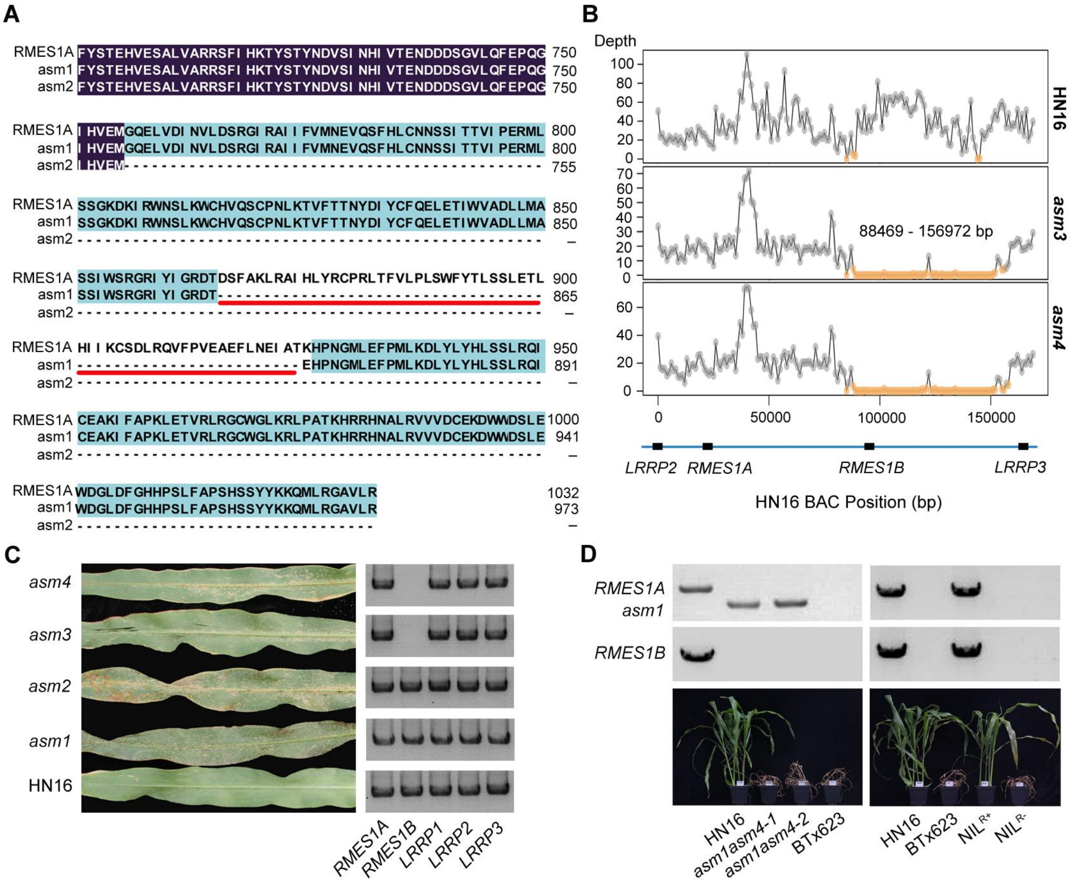

**Figure S1. Identification of mutants and near-isogenic lines, related to Figure 1.**

(A) Amino acid sequence comparison among RMES1A and two derivative mutants. While asm1 has an internal deletion of 59 amino acids, asm2 is prematurely terminated and lacked the deduced 277 aas in the C-terminus. The 59 residues deleted in asm1 are underlined.

(B) The deletions in *ams3* and *ams4* mutants were firstly analyzed using their genome resequencing reads, which were mapped to the 169 kb *RMES1*-carrying BAC contig (shown in blue), with the region showing low read coverage identified (marked in orange). Both mutants suffered a deletion of 68.503 kb, resulting in the loss of *RMES1B*.

(C) Reaction phenotypes to *MES* infestations among HN16 and four *asm* mutants. When grown in the field, abundant *MES* aphids were found on the leaves of the four mutants but not HN16. Genomic PCR assays show that *RMES1B* is completely missing in *asm3* and *asm4*. *LRRP1*, *LRRP2*, and *LRRP3* are the three paralogs of *RMES1A*/*RMES1B* in the *RMES1* genomic region.

(D) Identification of double mutant of *RMES1A* and *RMES1B* (left panel) and near-isogenic lines (NILs) with or without *RMES1A* and *RMES1B* (right panel). Genomic PCR assays were employed to check the presence or absence of *asm1, RMES1A*, and *RMES1B* in the relevant lines. The two sibling double mutant lines (*asm1asm4-1* and *-2*) and the NIL without *RMES1A* and *RMES1B* (NIL$^{R-}$) were susceptible to *MES* infestation, whereas the NIL with *RMES1A* and *RMES1B* (NIL$^{R+}$) showed resistance to *MES*. HN16 and BTx623 were used as *MES* resistant or susceptible controls. The graph was taken at 10 days post aphid inoculation.

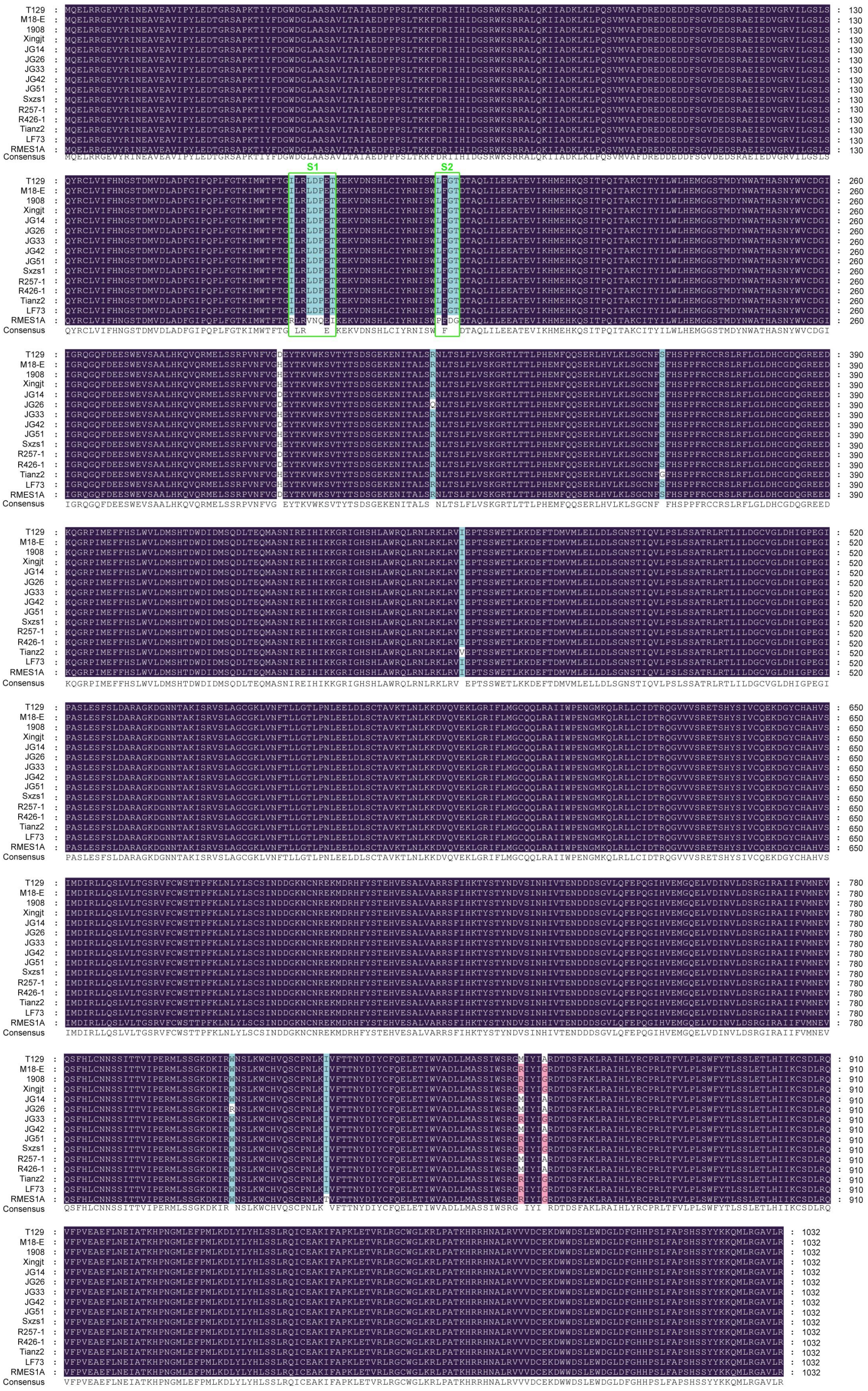

**Figure S2. Comparison of deduced amino acid sequences of *RMES1A* alleles in representative sorghum germplasm lines, related to Figure 1.**

Amino acid sequence alignment of *RMES1A* (from HN16) and its alleles in 14 susceptible sorghum lines (listed in Table S1). The positions with amino acid substitutions are highlighted by different colors. The main variable regions S1 (ILRLDPET) and S2 (LFGT) are indicated by green boxes.

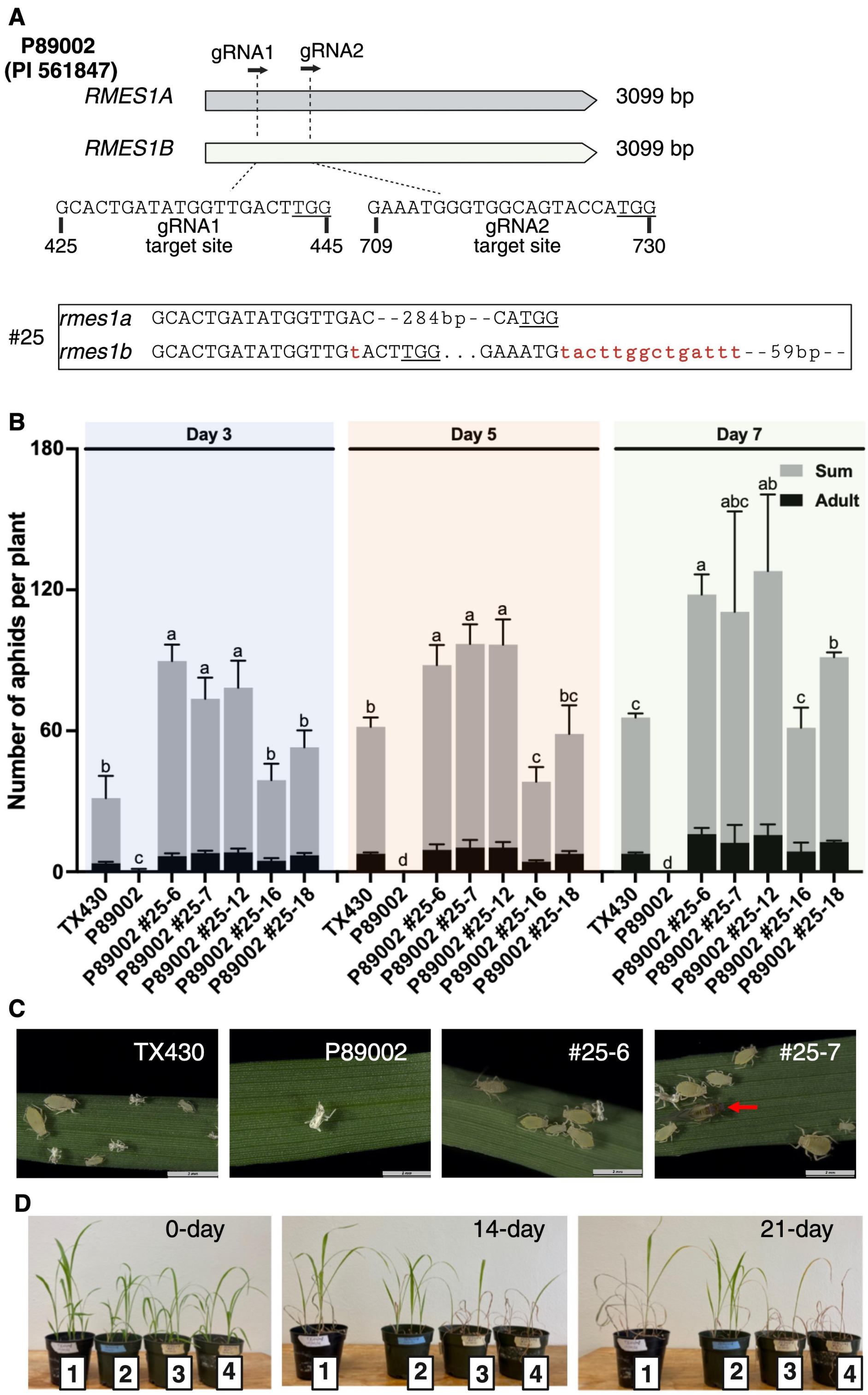


**Figure S3. Generation and assessment of a CRISPR mutant of *RMES1A* and *RMES1B*, related to Figure 1.**

(A) Coding regions of *RMES1A* and *RMES1B* and CRISPR-induced mutations in them. The *MES* resistant sorghum line P89002 (PI 561847), containing both *RMES1A* and *RMES1B*, was subjected to CRISPR/Cas9-mediated editing using gRNA1 (425 - 445 bp) and gRNA2 (709 - 730 bp). The two target sites were both conserved in *RMES1A* and *RMES1B*. One homozygous stable mutant (P89002 #25), which carried frameshift mutations in both genes, was obtained for further analysis. The dashed lines, lower case letters, and dots indicate deletions, insertions, and the bases not shown, respectively. The underlined nucleotides mark proto-adjacent motifs.

(B) Comparison of *MES* aphid numbers on different sorghum lines at 3-, 5- and 7-days post inoculation. TX430, P89002, and five familial lines derived from P89002 #25 were inoculated with adult aphids (5 plants per line with 10 aphids per plant in each experiment). Both adults and nymphs were counted at the indicated time points, with sum including adults and nymphs. Error bars, means ± SE (n = 3). Different letters above the histograms indicate significant differences by Tukey's HSD test ($p < 0.05$).

(C) Population status of *MES* feeding on different sorghum genotypes for 7 days. TX430 sustained a substantial *MES* population with newly produced nymphs, whereas most aphids died, and no nymphs were produced in P89002. The two CRISPR lines (P89002 #25-6 and #25-7) supported thriving *MES* populations, including alate *MES* aphids due to overcrowded colonies. The red arrow indicates an alate adult.

(D) Growth states of sorghum plants of different genotypes after being inoculated with a mixture of 100 nymphs and apterous adult aphids for 14 and 21 days. The genotypes of the plants are: 1, TX430; 2, P89002; 3, P89002 #25-6; and 4, P89002 #25-7. Symptoms of wilting and drooping were observed at 14 days post inoculation in the control TX430 group and in the CRIPSR lines #25-6 and #25-7. By day 21 of infestation, most P89002 #25-6 and #25-7 plants were dead, while those of P89002 continued to grow.

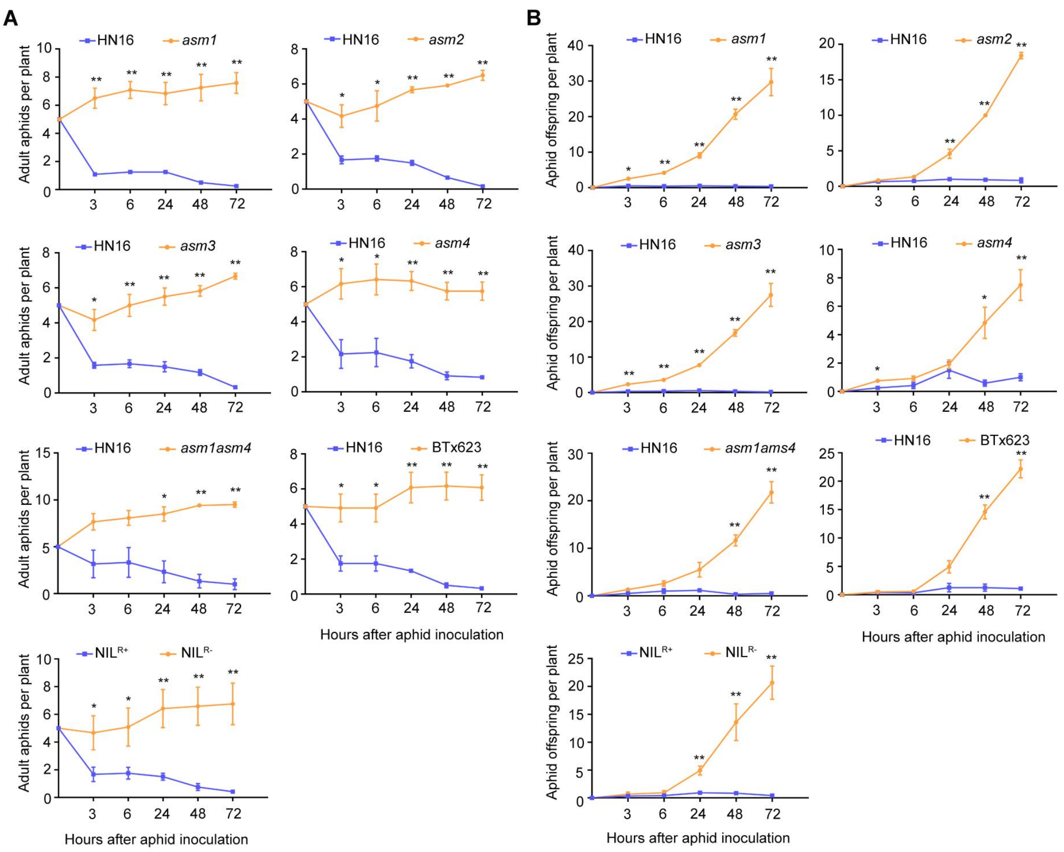


**Figure S4. Number of adults and nymphs produced by *MES* after feeding on different sorghum genotypes, related to Figure 2.**

(A) Settling of aphids on HN16, BTx623, two NILs, and five mutants in host choice tests. In each test, seven trials were conducted with each trial involving two genotypes. Five adult aphids were used for each genotype, and the number of aphids settled on individual genotypes was monitored at the indicated time points. Error bars, means ± SE (n = 3), $^{*}p < 0.05$, $^{**}p < 0.01$, Student's *t* test (compared to HN16 or $NIL^{R+}$).

(B) Number of nymphs produced by *MES* after inoculating on the leaves of different sorghum genotypes in no-choice tests. In each test, seven trials were conducted with each trial involving two genotypes. Ten adult aphids were used for each genotype, and the number of nymphs newly produced on individual genotypes was monitored at five time points. Error bars, means ± SE (n = 3), $^{*}p < 0.05$, $^{**}p < 0.01$, Student's *t* test (compared to HN16 or $NIL^{R+}$).

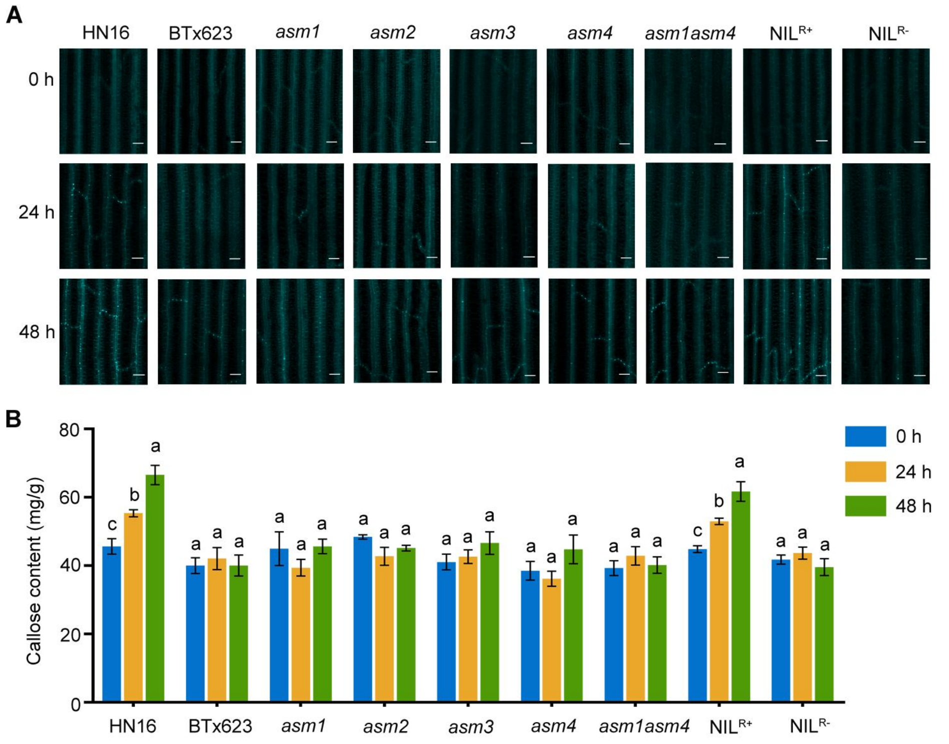


**Figure S5. *RMES1A* and *RMES1B* are required for stimulating callose deposition by *MES* feeding in sorghum, related to Figure 2.**

(A) Aniline blue staining of callose deposition in nine sorghum genotypes at 0, 24, and 48 h post *MES* inoculation. Stimulation of callose deposition (indicated by the bright dots around leaf veins) was observed in only HN16 and $NIL^{R+}$ that carry both *RMES1A* and *RMES1B*. Scale bars, 50 µm.

(B) Callose contents in the leaf tissues of nine sorghum genotypes at 0, 24, and 48 h post *MES* inoculation. Error bars, means ± SE (n = 3). Different letters above the histograms indicate significant differences by Tukey's HSD test ($p < 0.05$). Significant increase of callose content after *MES* inoculation was seen in only HN16 and $NIL^{R+}$ that carry both *RMES1A* and *RMES1B*.

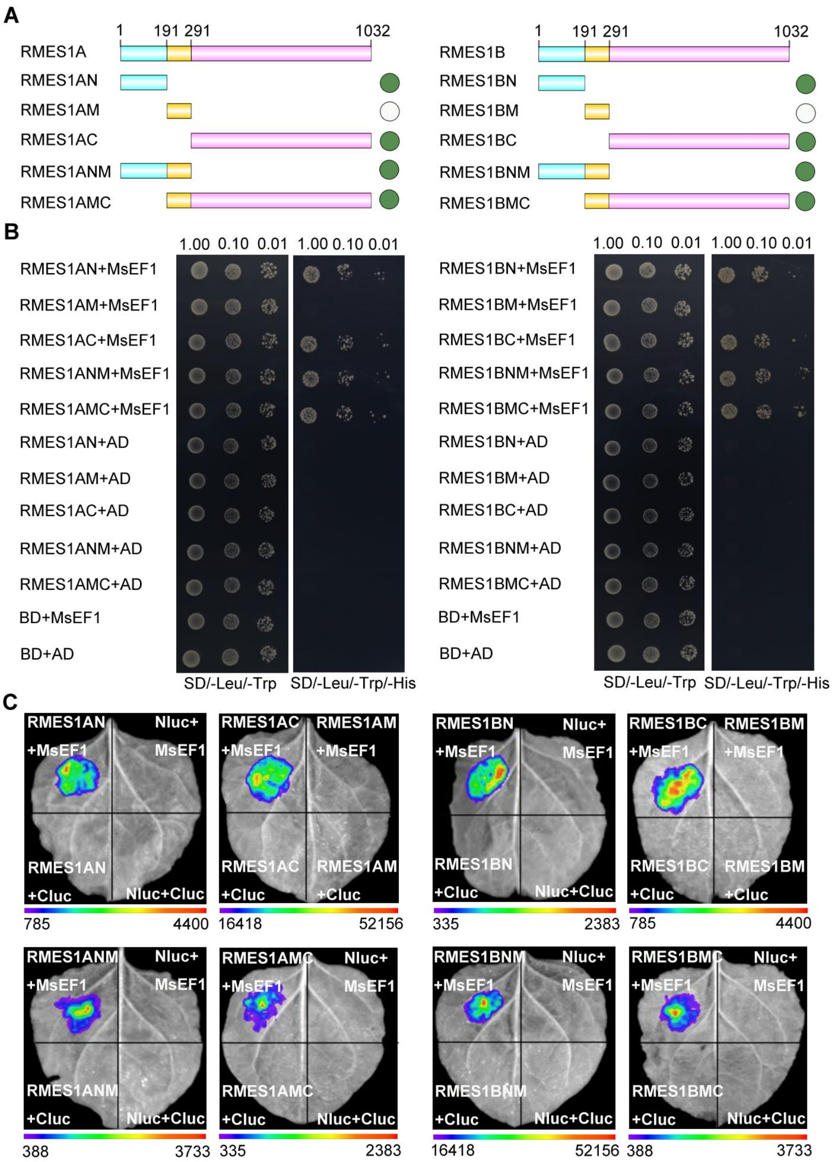


**Figure S6. Analysis of RMES1A and RMES1B regions involved in interacting with MsEF1, related to Figure 5.**

(A) A diagram showing the domain structure of RMES1A and RMES1B predicted using the AlphaFold 2 system (see Figure 7). N, M, and C indicate the putative N-terminus, middle domain, and C-terminus, respectively. For both RMES1A and RMES1B, five derivative fragments were examined for potential interaction with MsEF1 using yeast two hybrid (Y2H) and split luciferase complementation (SLC) assays. The green and white circles denote positive or negative interactions with MsEF1 according to the results shown in (B) and (C).

(B) Y2H assay results of the interaction between RMES1A/RMES1B and MsEF1. MsEF1 was fused with the activation domain (AD), and RMES1A/RMES1B derivative fragments were each fused with DNA-binding domain (BD) for Y2H assays, with positive yeast colony growth on the SD/-Leu/-Trp/-His medium indicating occurrence of protein-protein interaction. The empty AD and BD vectors were used as negative controls.

(C) SLC assay results of the interaction between RMES1A/RMES1B and MsEF1. MsEF1 was fused with the C-terminus of luciferase (Cluc), and RMES1A/RMES1B derivative fragments were each fused with the N-terminus of luciferase (Nluc) for SLC assays. The empty Nluc and Cluc vectors were employed as negative controls.

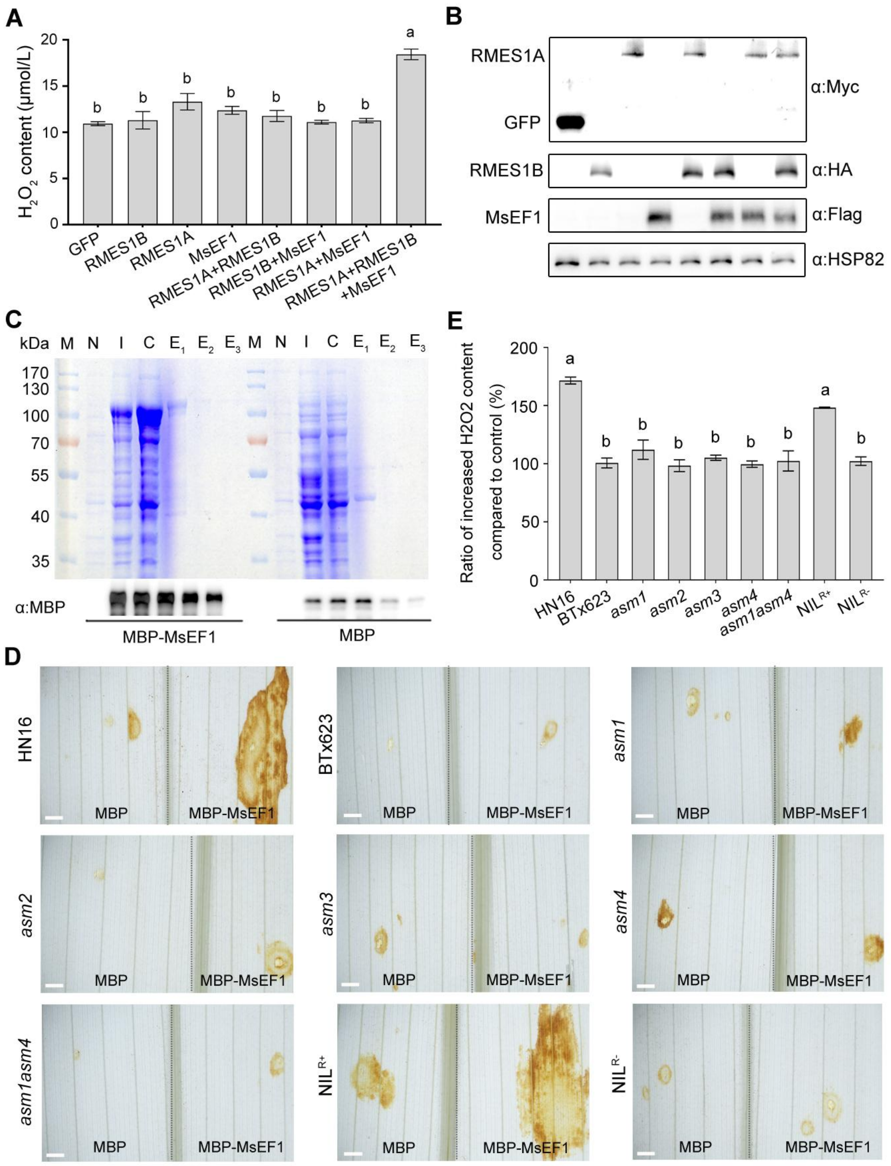


**Figure S7. MsEF1 elicits $H_2O_2$ burst in the presence of RMES1A and RMES1B, related to Figure 6.**

(A) MsEF1 upregulates $H_2O_2$ content in BTx623 protoplasts in the presence of ectopically expressed RMES1A and RMES1B. Flag-tagged MsEF1 was co-expressed with Myc-tagged RMES1A and/or HA-tagged RMES1B in the protoplasts, with $H_2O_2$ content measured at 24 h after protoplast transfection. Expression of GFP was used as a negative control. Error bars, means ± SE (n = 3). Different letters above the histograms indicate significant difference according to Turkey's HSD test ($p < 0.05$).

(B) Immunoblotting assays confirming the expression of Flag-tagged MsEF1, Myc-tagged RMES1A, HA-tagged RMES1B, and GFP in the transfected protoplast samples. Immuno-detection of HSP82 served as a loading control.

(C) Purification of MBP-MsEF1 and MBP recombinant proteins inducibly expressed in bacterial cells. The upper panel shows the result of 12% SDS-PAGE, separating protein molecular markers (M), total cell extract of the bacterial culture prior to IPTG induction (N), total cell extract after 0.1 mM IPTG induction (I), crude extract of lysed cells after induction (C), and recombinant proteins eluted from amylose column with 10 mM, 20 mM, or 50 mM maltose ($E_1$-$E_3$). The bottom panel depicts immunoblotting result of MBP-MsEF1 and MBP in the different steps of purification. MBP antibody was used in the immunoblotting assays.

(D) Elicitation of strong $H_2O_2$ burst by purified MBP-MsEF1 in the genotypes carrying both RMES1A and RMES1B (HN16 and $NIL^{R+}$) but not those with functional deficiency of RMES1A and/or RMES1B (BTx623, five asm1 mutants, and $NIL^{R-}$). Purified MBP was used as a negative control. The upper panel shows DAB staining result. For each genotype, equal amount (0.5 μg) of MBP or MBP-MsEF1 was pressure-injected into opposite areas of the same leaf, with DAB staining conducted at 36 h after the injection. Scale bars, 50 μm.

(E) Quantitative measurement result of $H_2O_2$ contents for the nine different treatments shown in (D). The ratio of increased $H_2O_2$ content was calculated by dividing the amount of $H_2O_2$ induced by MBP-MsEF1 with that by MBP. Error bars, means ± SE (n = 3). Different letters above the histograms denote significant difference determined by Tukey's HSD test ($p < 0.05$).

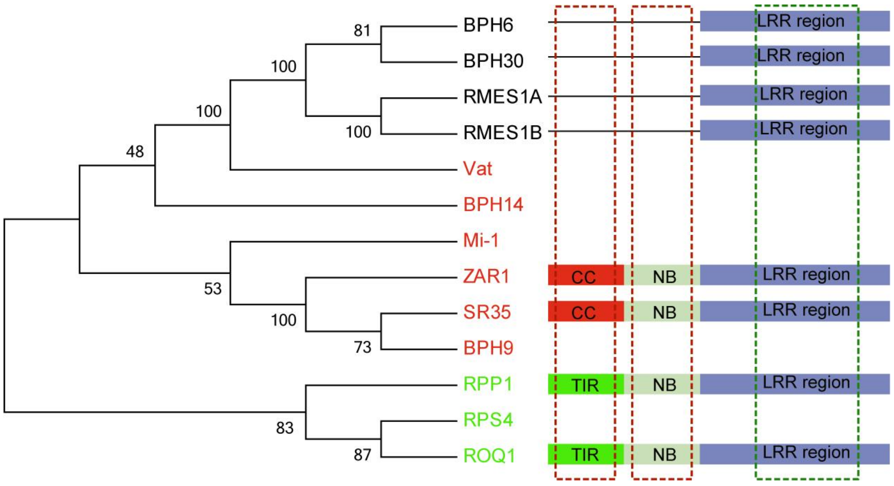


**Figure S8. Phylogenetic analysis of RMES1A and RMES1B and investigation of their potential domains through amino acid sequence comparison with typical CNLs and TNLs, related to Figure 7.**

Phylogenetic analysis shows that RMES1A, RMES1B, BPH6 and BPH30 form a distinct cluster separated from typical CNLs (represented by ZAR1, SR35 and previously cloned insect resistance proteins Mi-1, Vat, BPH9 and BPH14) and canonical TNLs (e.g., RPP1, RPS4 and ROQ1). RMES1A, RMES1B, BPH6 and BPH30 resemble CNLs and TNLs in having a leucine-rich repeat (LRR) region in the C-terminus. But no homologous coiled coil (CC), Toll/interleukin-1 receptor (TIR), and nucleotide binding (NB) domains could be detected in RMES1A, RMES1B, BPH6 and BPH30 through amino acid sequence comparison with CNLs and TNLs. The tree shown was constructed using the neighbor joining program installed in MEGA 11 (https://www.megasoftware.net/), with the bootstrap values obtained through 1000 permutations.

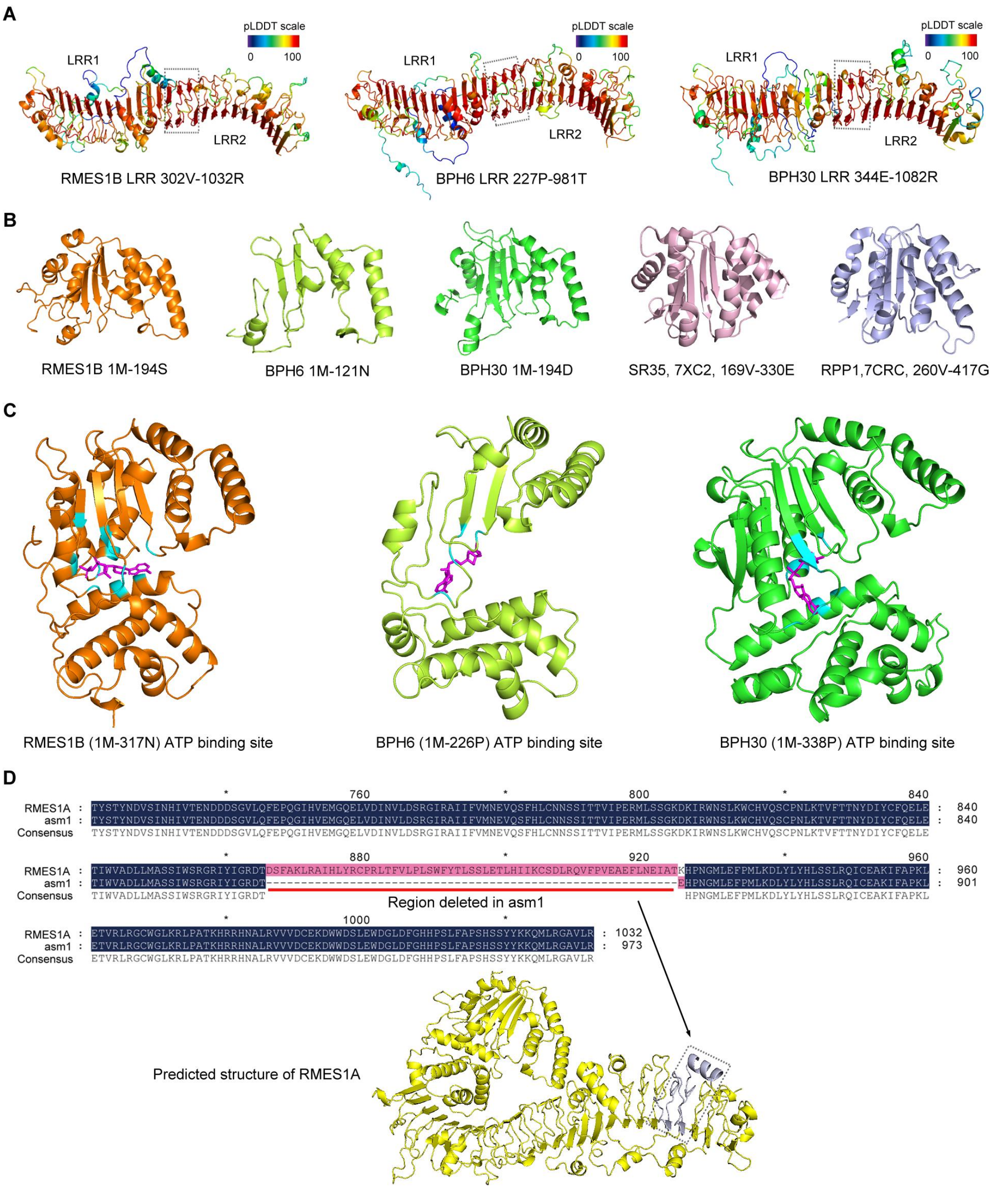


**Figure S9. Predicted structural features of RMES1B, BPH6, BPH30, and asm1, related to Figure 7.**

(A) Two putative LRR domains were predicted RMES1B, BPH6, and BPH30. The three pairs of anti-parallel β-strands separating the two LRR domains were boxed.

(B) The putative N-terminal domain of RMES1B, BPH6, and BPH30 resembles the nucleotide binding domain of SR35 and RPP1 in both structural composition (α-helices + β-strands) and overall topology.

(C) A potential ATP-binding site, located in a grove between the putative N-terminal and middle domains, is similarly predicted for RMES1B, BPH6, and BPH30. ATP is shown in purple, while the structural elements possibly involved in ATP binding are marked in light blue.

(D) Compared to wild type RMES1A, the asm1 mutant has an internal deletion of 59 amino acids in its C-terminal region. The 59 residues deleted in ams1 correspond to two LRRs in the second LRR domain of the predicted RMES1A structure.
